# Generative design of antigen-specific T-cell receptor sequences with a conditional diffusion model

**DOI:** 10.64898/2026.06.10.730756

**Authors:** Yumeng Zhang, Wenhua Liang, Shuting Xu, Matthew Witney, Xiao-Dong Su, Miles C. Andrews, Jamie Rossjohn, Anthony W. Purcell, Feng Wang, Jiangning Song

## Abstract

T cell receptor (TCR)-based immunotherapy holds immense potential for treating cancers, autoimmunity, and infectious diseases, where antigen-specific TCR recognition is crucial for adaptive immune responses. Engineering or *de novo* generation of the complementarity-determining region 3 (CDR3) loops of TCRs using artificial intelligence offers a powerful alternative to designing antigen-specific TCRs rather than laborious experimental screening. However, current *in silico* approaches are constrained by weak conditional guidance, limited flexibility, and a lack of rigorous functional validation. To address these limitations, we introduce TCRDiff, a generative diffusion framework for designing antigen-specific TCRs conditioned on peptide-MHC (pMHC) targets and germline-encoded TCR variable genes. By leveraging pre-trained knowledge from massive T-cell repertoires and TCR-pMHC recognition data, TCRDiff generates CDR3αβ sequences that closely resemble native-binding TCRs via a denoising diffusion process. Furthermore, incorporating interface geometry features generated TCR-pMHC complexes with superior structural plausibility than models relying solely on sequence-based diffusion or structure-based modeling. As a proof of concept, we deployed TCRDiff in a systematic pipeline to design candidate TCRs against a clinically validated cancer antigen. *In vitro* activation assays validated that TCRDiff-generated TCRs efficiently recognize the MAGE-A3 epitope with minimal off-target reactivity. Thus, TCRDiff establishes a powerful, validated computational paradigm to accelerate the development of TCR-based immunotherapies.

## Main

T lymphocytes are central to adaptive immunity, playing a pivotal role in combating viral infections and cancers, while their dysregulation can drive autoimmune disorders^1^. T cell-mediated immunity is primarily governed by the molecular interactions between paired T cell receptor (TCR) *αβ* chains and antigen-derived peptides presented by major histocompatibility complex (MHC) molecules^2^, which trigger T-cell activation and orchestrate downstream immune responses. Critically, the hypervariable complementary-determining region 3 (CDR3) loops often contribute substantially to peptide recognition and T-cell specificity, while CDR1 and CDR2 loops can also mediate important contacts with both peptide and MHC^3^. Concurrently, the conformational flexibility of the CDR loops may allow a single TCR to accommodate distinct antigens, thereby contributing in part to the structural basis for cross-reactivity^4^.

TCRs demonstrate increasing therapeutic potential in cancer immunotherapy^5^, notably in tumor-infiltrating lymphocyte (TIL) therapy^6, 7^, TCR-based bispecific T-cell engager (BiTE)^8^, and TCR-engineered T (TCR-T) cell therapy^9^. A crucial step in developing safe and effective immunotherapies is to identify antigen-specific TCRs. Specifically, TIL therapy requires isolating tumor-reactive T cells directly from patient samples^10^, while TCR-based BiTE and TCR-T cell therapies typically rely on engineering TCRs that target known antigens^11^. Currently, the experimental discovery of antigen-specific TCRs relies heavily on peptide-MHC (pMHC)-based high-throughput screening methods, such as MHC tetramer-associated TCR sequencing (TetTCR-seq)^12, 13^, microfluidic antigen-TCR engagement sequencing (MATE-seq)^14^, and barcode-enabled antigen mapping of T cells (BEAM-T)^15^. Additionally, phage and yeast display systems^16–19^ enable the directed evolution of TCR mutants to improve binding affinity and specificity. However, existing experimental techniques face critical bottlenecks. Tumor-reactive T cells are often rare in peripheral blood and difficult to detect due to the inherently low-affinity of the TCR-pMHC interaction^20^. Furthermore, current experimental platforms require labor-intensive T-cell isolation and are not designed to efficiently explore the vast TCR sequence space while simultaneously optimizing antigen specificity and minimizing off-target reactivity. Consequently, extensive downstream functional assays remain indispensable for validating the cytotoxicity and potency of candidate T cells, as well as for reducing off-target effects^21^. This creates a significant hurdle in the development of safe and effective TCR-based therapeutics. Therefore, a pressing need exists for computational frameworks capable of designing or prioritizing antigen-specific TCRs with improved specificity and functional potential, thereby accelerating therapeutic development^22^.

Current deep learning-based generative models offer a promising direction for designing antigen-specific TCR sequences; however, these approaches still face major limitations. For instance, while TCR-VALID^23^ transforms TCR sequences into a low-dimensional disentangled space using C*β*-Variational Autoencoder (VAE), this antigen-agnostic model decodes meta-representations of TCR clusters rather than generating antigen-specific TCR sequences. Models like ERTransformer^24^, GRATCR^25^, and TCR-Translate^26^ employ sequence-to-sequence (Seq2Seq) frameworks^27^ for *de novo* TCR sequence generation directly from peptide or peptide-MHC complex (pMHC) inputs. However, these frameworks are restricted to designing CDR3*β* sequences and to antigenic peptides presented by human MHC-I molecules. As germline-encoded V and J genes constrain the conserved regions within the CDR3 loop, TAPIR^28^ supports the *de novo* design of paired TCR sequences by sequentially predicting both CDR3*α* and CDR3*β* given a target pMHC and VJ genes. Nevertheless, such generation strategies still suffer from weak conditional guidance and limited flexibility to optimize key peptide-interacting sites along the CDR3αβ loops. In contrast, discrete diffusion models^29–31^ offer stronger conditional controllability and iterative refinement, thereby enabling more flexible and precise generation of antigen-specific TCRs. Finally, structure-based sequence design methods, such as ProteinMPNN^32^, often fail to generalize to specific interaction conformations between peptides and CDR3 loops due to the limited availability of high-resolution crystal structures of TCR-pMHC complexes^33^.

Another category of methods focuses on developing chimeric antigen receptor (CAR)-like molecules binding to pMHC targets^34–39^. Combinatorial library screening guided by molecular docking, or computational design pipelines incorporating RFdiffusion^40^, ProteinMPNN^32^, and AlphaFold^41^, have successfully designed functional pMHC binders for classic tumor-associated antigens such as NY-ESO-1^42^. While these strategies demonstrate promising therapeutic potential, their objective fundamentally differs from designing antigen-specific TCR sequences. Engineered small-molecule pMHC binders are designed for high-affinity, peptide-specific recognition that can potentially operate across divergent MHC contexts. In contrast, TCR-based designs leverage the natural structural scaffold of TCRs to recognize intracellular tumor antigens. This preserves better compatibility with canonical pMHC docking geometry, ensuring proper TCR-mediated signaling and downstream T-cell activation^43^.

Here, we present TCRDiff, a conditional discrete diffusion model for the *de novo* generation of antigen-specific TCR CDR3αβ sequences. By training on cross-organism and pan-MHC TCR-pMHC reactive pairs, TCRDiff demonstrates enhanced generalizability in predicting TCR binding specificity. During the denoising diffusion process, conditional guidance from both the encoded pMHC target and the TCR context enables iterative generation of amino acids within the CDR3αβ loops. TCRDiff achieves state-of-the-art conditional generation performance, as evaluated by both sequence consistency via deterministic decoding and CDR3 motifs via stochastic generation. Furthermore, this flexible generation framework can integrate knowledge of the interface geometry constraints derived from structure-based sequence design methods, thereby yielding designed TCR-pMHC complexes with superior structural plausibility. To streamline candidate selection for immunotherapy, we establish a pipeline combining TCR generation, *in silico* screening, and *in vitro* functional validation. We successfully apply this pipeline to design highly specific TCRs targeting the MAGE-A3^44, 45^ epitope presented by the HLA-A*01:01 molecule with limited off-target reactivity observed against a known self-peptide. We envision TCRDiff as a powerful computational framework that will contribute to the development of safe and effective TCR-based immunotherapies.

## Results

### Overview of TCRDiff

TCRDiff is a pMHC-conditioned discrete diffusion language model of TCR CDR3αβ sequences (**Fig. 1a**). At each diffusion timestep, TCRDiff learns to denoise masked amino acid tokens using partial CDR3αβ sequences combined with conditional knowledge of pMHC binding specificity, thereby reconstructing complete CDR3 sequences. The network architecture of TCRDiff is illustrated in **Fig. 1b** and **Supplementary Fig. 1**. The input tokens of paired CDR3αβ sequences are first processed by Diffusion Transformer (DiT) blocks^46^ to capture the biological grammar of CDR3 loops constrained by germline-encoded V genes. This unsupervised DiT-based language model was pre-trained on TCR sequences extracted from human and mouse T-cell repertoires (Refer to the “Methods” section for details). Meanwhile, the target peptide sequence and MHC molecule are processed by a peptide-MHC binding model, which was trained to quantify pMHC binding affinity. The single and pairwise representations derived from the model’s intermediate PairFormer block^41^ characterize the interaction patterns within the pMHC complex. A PairFormer adapter then refines the TCR representations by integrating their interaction potential with specific pMHC targets. Finally, the complete CDR3αβ sequences are decoded by the language model head during the denoising diffusion process (Refer to the “Methods” section for details). To ensure that the conditional diffusion model accurately drives the generation of antigen-specific TCRs, TCRDiff leverages the pre-trained weights of a TCR-pMHC binding predictor built on the same architecture. They were trained on a diverse dataset of TCR-pMHC reactive pairs, spanning human and mouse TCRs, as well as MHC class I and II peptides.

**Fig. 1.**
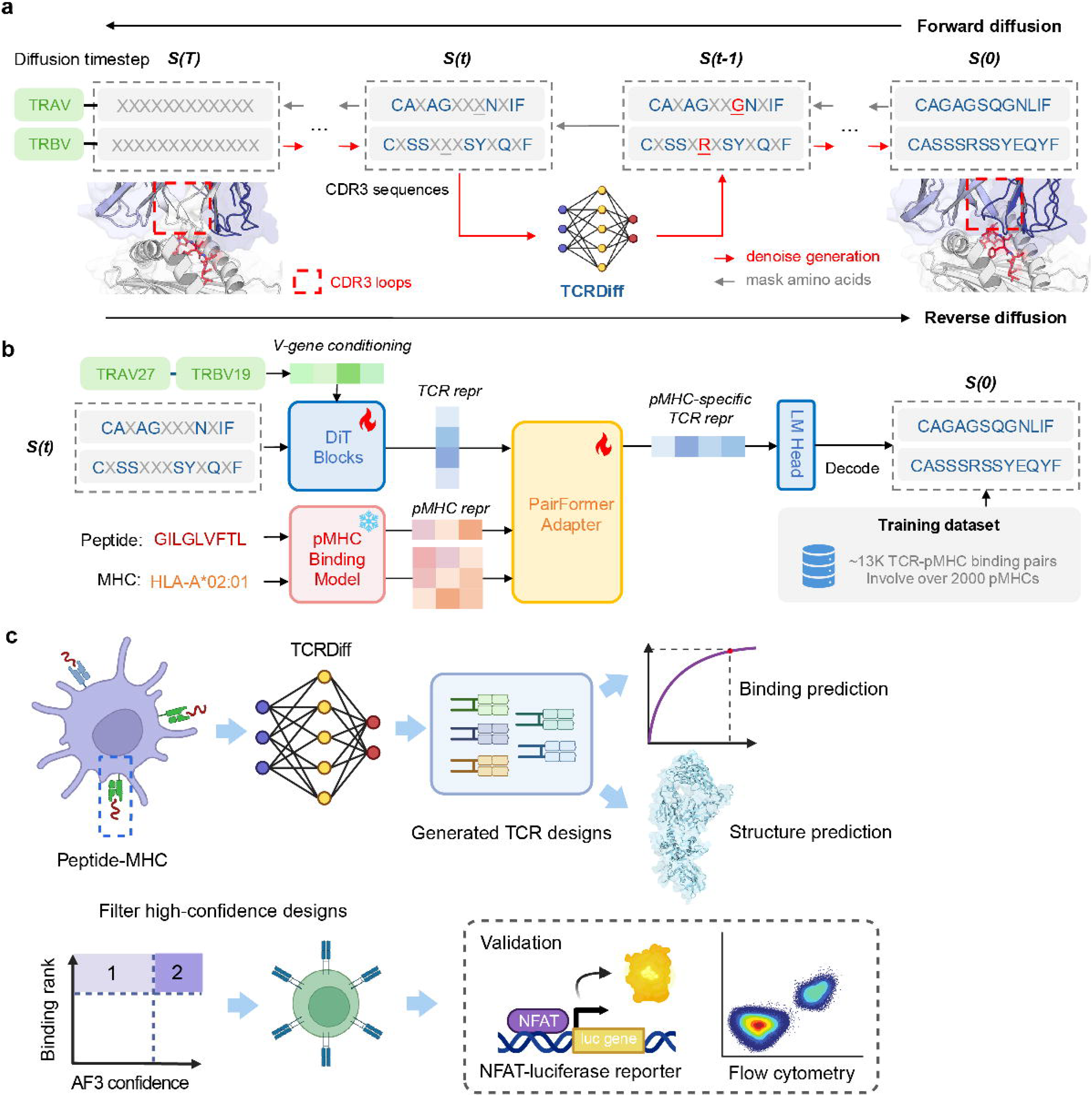
TCRDiff: a pMHC-conditioned diffusion model for de novo T-cell receptor design. **a**, Schematic diagram of the diffusion process for TCR CDR3αβ sequences. During the forward diffusion phase, native CDR3αβ sequences are incrementally masked; TCRDiff is employed to iteratively denoise the masked CDR3αβ sequences during the reverse diffusion process. **b**, Architecture of the TCRDiff model, which comprises a pre-trained pMHC binding model, trainable Diffusion Transformer (DiT) blocks and a PairFormer adapter to reconstruct the original CDR3αβ sequences from the reactive TCR-pMHC pairs. **c**, Schematic of the TCR design pipeline, which incorporates *in silico* conditional generation, multi-tiered structural and binding filtering, and downstream *in vitro* functional validation of TCRs tailored to specific peptide-MHC targets.

As TCRDiff generates potential reactive TCRs for pMHC targets of interest, we establish an antigen-specific TCR design pipeline (**Fig. 1c**). The sequences generated by TCRDiff effectively capture the specific amino acid preferences of CDR3αβ sequences across distinct tumor or viral antigens. Following sequence generation, TCR-pMHC binding predictors and structural modeling frameworks for protein complexes, such as AlphaFold, enable effective filtering and prioritization of highly confident TCR candidates based on binding ranks and structural confidence metrics^47^. These prioritized TCR designs are introduced into T cells, and *in vitro* functional assays are subsequently performed to validate their reactivity against specific target epitopes.

### TCRDiff achieves state-of-the-art performance in predicting TCR-pMHC reactive pairs

Building on the divide-and-conquer workflow of our previous TCR-pMHC binding predictor^48^, we expanded the data sources and upgraded the model architecture for a more comprehensive understanding of TCR-pMHC binding specificity (**Fig. 2a**). As a unified predictor, TCRDiff expands beyond human TCR*αβ*-pMHC-I pairs to also accommodate MHC class II molecules, single-chain TCRs, and mouse data inputs. The model implements a combined sampling strategy to generate “non-binding” TCRs^49^, performs a two-stage pretraining^31^ on unlabeled TCR sequences using DiT, and utilizes a PairFormer module to capture residue-level interactions across both peptide-MHC and TCR-pMHC (See the “Methods” section for details).

**Fig. 2.**
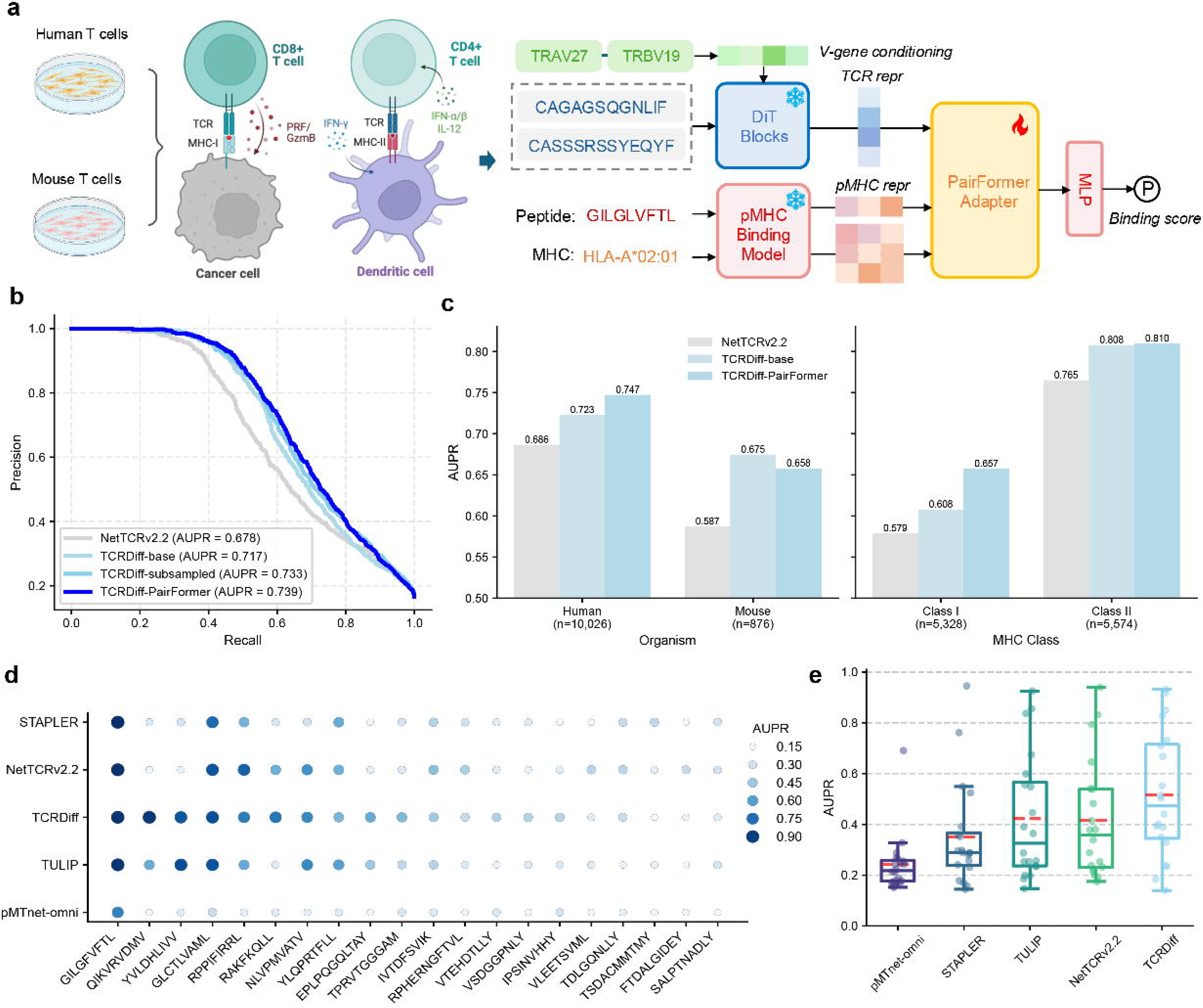
TCRDiff demonstrates robust generalizability in predicting TCR-pMHC binding specificity. **a**, Schematic overview of the TCRDiff binding model. TCRDiff incorporates paired human and mouse TCR sequences and pan-MHC data into a unified model to predict TCR-pMHC binding specificity. **b**, Precision-recall curves comparing NetTCR v2.2 against different versions of TCRDiff evaluated on the validation dataset. **c**, Bar plots displaying AUPRs achieved by NetTCR v2.2, TCRDiff-base, and TCRDiff-PairFormer, stratified by organisms (left) and MHC classes (right). **d**, Dot plots displaying AUPRs achieved by pMTnet-omni, STAPLER, TULIP, NetTCR v2.2, and TCRDiff across individual peptides in the IMMREP23 test set. Larger size and darker color of the point indicate a higher AUPR value. **e**, Box plots displaying the distribution of AUPRs across all peptides in the IMMREP23 test set. Box center line, median; dashed line, mean; box limits, upper and lower quartiles; whiskers, 1.5 ×interquartile range; points, data points.

We first established two baseline models for performance evaluation: NetTCR v2.2^50^, which directly concatenated sequence features of the six TCR CDR loops, and TCRDiff-base, which utilized the same contrastive learning approach as EPACT^48^. To mitigate performance bias driven by the most dominant pMHC targets, we also trained TCRDiff on a subsampled dataset, yielding TCRDiff-subsampled and TCRDiff-PairFormer. Ablation studies on the validation set demonstrated the incremental benefits of our design, as the overall area under the precision-recall curve (AUPR) increased sequentially with the introduction of pre-training (from 0.678 to 0.717), subsampling (from 0.717 to 0.733), and the PairFormer-based architecture (from 0.733 to 0.739) (**Fig. 2b**). Fine-tuning the last pre-trained DiT block yielded the top validation AUPR of 0.750 (**Extended Data Fig. 1a**). TCRDiff consistently outperformed NetTCR v2.2 across different organisms and MHC classes (**Fig. 2c**). TCRDiff-PairFormer achieved substantial improvements of AUPR over TCRDiff-base on both the human (0.747 vs. 0.723) and MHC-I (0.657 vs. 0.608) subsets, while maintaining comparable performance on the mouse (0.658 vs. 0.675) and MHC-II (0.810 vs. 0.808) subsets. When performance was stratified across MHC allotypes with at least 10 TCR-pMHC reactive pairs in the validation set, TCRDiff-PairFormer outperformed the other two baseline models in 10 out of 19 allotypes (**Extended Data Fig. 1b**). Notably, although MHC class I and II molecules present peptides in distinct modes with different TCR docking geometries^51^ during antigen recognition, the pan-MHC TCRDiff-PairFormer model achieved equivalent performance to the class-specific models (**Supplementary Fig. 2**).

We further evaluated TCRDiff on the private test set from the IMMREP23 TCR-epitope prediction challenge^52^, which comprises paired reactive TCRs for 20 pHLA-I targets. On this independent dataset, TCRDiff-PairFormer still outperformed TCRDiff-base and TCRDiff-subsampled, with the overall AUPR rising from 0.572 to 0.614 (**Extended Data Fig. 1c**). This performance gain highlights the robust generalizability empowered by the PairFormer architecture. Furthermore, TCRDiff exhibited remarkable performance improvements compared to existing TCR-pMHC binding predictors, including NetTCR v2.2^50^, STAPLER^53^, and pMnet-omni^54^ on the private benchmarking dataset (**Extended Data Fig. 1d**). When benchmarking model performance for individual peptide targets, TCRDiff achieved the best among five computational methods for 8 out of 20 peptides (**Fig. 2d**). Notably, the model substantially boosted the predictive accuracies for several critical viral antigens, including CMV epitopes: QIKVRVDMV (AUPR=0.918), TPRVTGGGAM (AUPR=0.504), and VTEHDTLLY (AUPR=0.402), EBV epitopes: RAKFKQLL (AUPR=0.732) and EPLPQGQLTAY (AUPR=0.552), and influenza epitope: VSDGGPNLY (AUPR=0.393). The results represent a 23.8% to 67.5% performance improvement over the second-best method for these targets. Generally, TCRDiff achieved state-of-the-art performance when evaluated across the average metrics for 20 peptides (**Fig. 2e**), yielding an average AUPR of 0.515, compared to the second-best model, TULIP^55^, which achieved an average AUPR of 0.422. To control the false positive rate (FPR) and emphasize high sensitivity, we also analyzed the area under the ROC curve up to an FPR of 0.1 (AUC0.1) (**Extended Data Fig. 1e, f**). TCRDiff maintained a decisive competitive advantage, achieving an average AUC0.1 score of 0.665 compared to TULIP’s AUC0.1 score of 0.620.

Identifying functional TCRs specific to cancer neoantigens, especially noncanonical antigenic peptides, holds immense therapeutic potential for cancer immunotherapy. To assess the generalizability of TCRDiff to neoepitopes distinct from those in the training set, we evaluated its performance on four cryptic peptides restricted to either HLA-A*02:01 or HLA-A*11:01^56^, alongside their reactive TCRs identified via a BEAM-T workflow. We calculated the binding rank percentiles of TCRs against a background control of 10,000 TCRs sampled from healthy human T-cell repertoires. For three out of four noncanonical pHLA targets (NU11, NU42, NU46, and NU49 antigens), more than half of the reactive TCRs ranked within the top 10% of the model’s prediction score distributions (**Supplementary Fig. 3**). Notably, the predictive performance was markedly better for peptides presented by HLA-A*02:01, where reactive TCRs ranked within the top 3%, compared to those presented by the HLA-A*11:01 allotype.

**Fig. 3.**
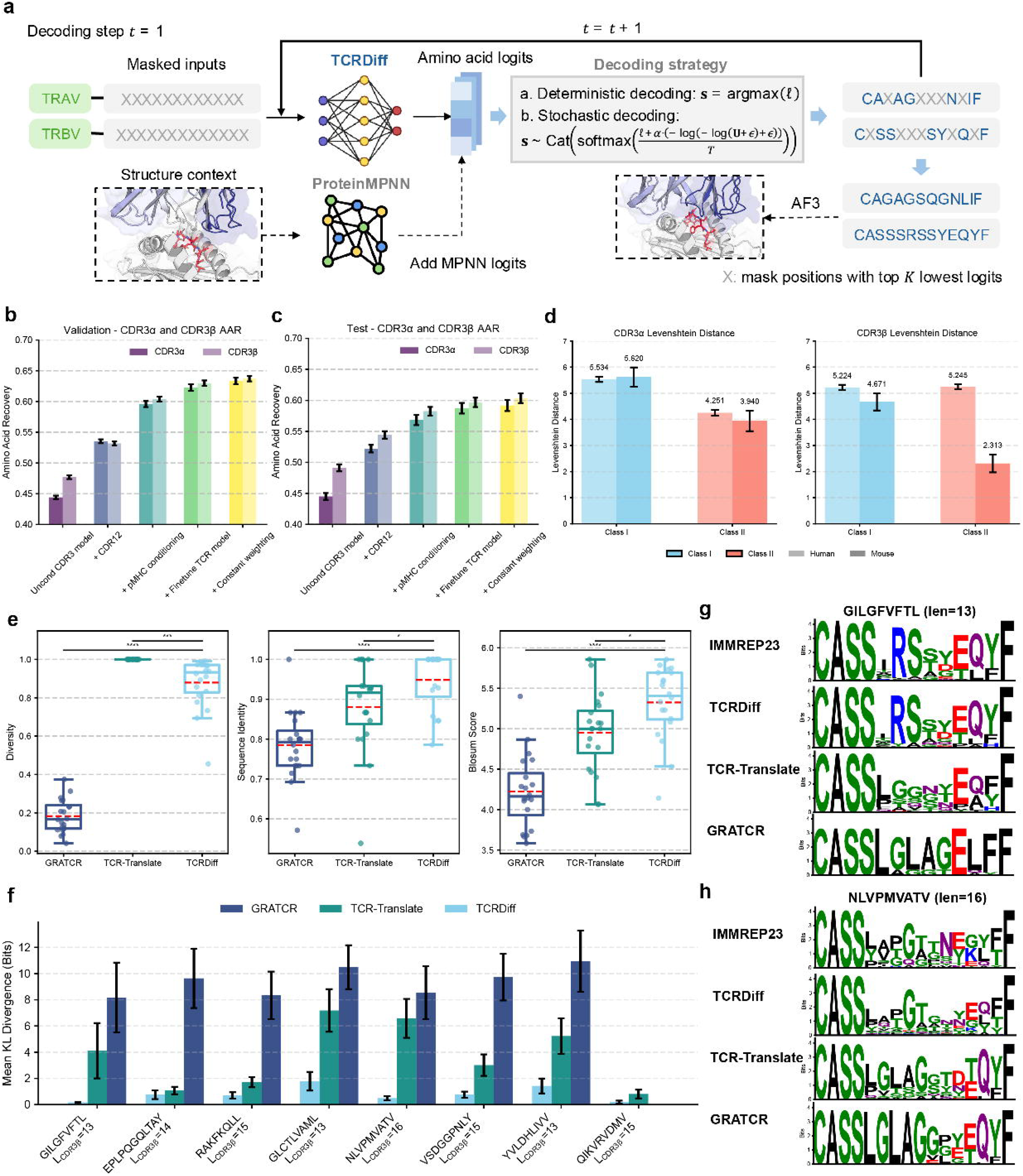
TCRDiff enables highly consistent CDR3 sequence generations compared to native sequences. **a**, Schematic overview of the iterative generation process of CDR3αβ sequences by TCRDiff. The workflow includes deterministic and stochastic decoding strategies, alongside the integration of optional structural conditions via ProteinMPNN’s predicted logits (TCRDiff+). **b, c**, Ablation study of deterministic generation by TCRDiff across various conditional inputs and training paradigms, evaluated on the validation set (b) and the IMMREP23 test set (c). Bar, AAR for CDR3*α* and CDR3*β*; error bar, standard error of AAR. **d**, Bar plots displaying the Levenshtein distances between the generated TCRs and native CDR3*α* (left) and CDR3*β* (right) sequences, stratified by organisms and MHC classes. Error bar, standard error of the Levenshtein distance. **e**, Benchmarking of stochastic generation performance (left, diversity; middle, sequence identity; right, BLOSUM score) among GRATCR, TCR-Translate, and TCRDiff on the IMMREP23 test set. Box center line, median; dashed line, mean; box limits, upper and lower quartiles; whiskers, 1.5×interquartile range; points, data points. Student’s t-test P values: *, P<0.05; **, P<0.01; ***, P<0.001; ****, P<0.0001. **f**, Mean Kullback-Leibler (KL) divergences of CDR3*β* motifs generated by GRATCR, TCR-Translate, and TCRDiff across diverse target peptides compared to the ground-truth motif derived from the IMMREP23 test set. Error bar, standard error of the KL divergence. **g, h**, Comparison of the sequence logos of CDR3*β* motifs generated by GRATCR, TCR-Translate, TCRDiff, and from the IMMREP23 test set for the peptides GILGFVFTL (g) and NLVPMVATV (h), fixed at the most frequent length of the CDR3*β* loop. The height of each amino acid letter is proportional to its sequence information content, measured in bits.

### TCRDiff generates antigen-specific TCRs for cross-organism and pan-MHC targets

To design TCR CDR3αβ sequences conditioned on pMHC binding specifications, TCRDiff employs an iterative generation procedure consisting of sequential decoding and re-masking operations throughout the denoising diffusion process (**Fig. 3a**). Specifically, TCRDiff predicts position-specific amino acid probability distributions within the CDR3αβ loops via two strategies: deterministic decoding, which selects the amino acid token with the maximum logit value, and stochastic decoding, which samples amino acid tokens based on the perturbed logits by Gumbel noise^57^.

We first evaluated the deterministic generation performance of TCRDiff on both the validation set and the IMMREP23 test set. TCRDiff recovered larger proportions of the fully masked CDR3αβ sequences by progressively integrating multi-aspect conditional information (**Fig. 3b, c**). The unconditional generation model accurately predicted approximately 46% of the amino acids within CDR3αβ sequences, a baseline primarily driven by the conserved regions at the N- and C-termini. Incorporating germline-encoded CDR1 and CDR2 features reinforced the model’s capacity to capture these conserved patterns, increasing the Amino Acid Recovery rates (AARs) to 0.534 on the validation set and 0.533 on the IMMREP23 test sets, respectively. As antigen recognition is predominantly mediated by the hypervariable core regions of CDR3 loops, integrating the pMHC binding conditions via the PairFormer adapter substantially boosted the generation accuracy, yielding AARs of 0.600 and 0.575 on the validation and test set, respectively. The final optimized TCRDiff model, which leveraged fine-tuning and a constant weighting of diffusion loss, achieved peak AARs of 0.635 and 0.597 (**Extended Data Fig. 2a, b**).

Stratifying the validation dataset by organism and MHC allotypes revealed that TCRDiff achieved strong generative performance on MHC class II data, characterized by higher AARs and shorter Levenshtein distances to native binding TCRs (**Fig. 3d** and **Extended Data Fig. 2c**). Interestingly, TCRDiff-generated mouse CDR3*β* sequences exhibited higher sequence similarity to natural counterparts than human CDR*β* sequences, particularly for pMHC-II targets (mean Levenshtein distance: 2.313 vs. 5.245). This discrepancy is likely attributable to the restricted diversity of mouse TCR repertoires within the training data. Finally, the average CDR3αβ AARs for the most frequent MHC allotypes in the validation set and the 20 peptides in the IMMREP23 test set (**Extended Data Fig. 2d, e**) demonstrated strong positive correlations with the AUPRs derived from the binding predictor (Spearman correlation coefficients of 0.614 and 0.738, respectively).

To evaluate stochastic generation, we generated 10 times the size of the native TCR-pMHC reactive pairs in the IMMREP23 dataset and subsequently extracted the top-ranked design for each pMHC target. Approximately 87.9% of the TCRDiff-generated CDR3*β* sequences were unique, demonstrating a sequence diversity comparable to that of TCR-Translate’s generations^26^. In contrast, another sequence-to-sequence model, GRATCR^25^, exhibited a highly restricted sequence diversity of only 18.3%. Furthermore, TCR designs generated by TCRDiff demonstrated superior maximum sequence consistency with the native TCRs in the IMMREP23 dataset (**Fig. 3e**). Specifically, TCRDiff achieved an average sequence identity of 94.8% and an average BLOSUM score of 5.33. These values were significantly higher than those achieved by GRATCR (78.5%, P=1 × 10^-7^; 4.22, P=3 × 10^-9^) and TCR-Translate (88.1%, P=0.016; 4.95, P=0.027). Whereas most existing methods focus primarily on CDR3*β* modeling, TCRDiff additionally supports the generation of antigen-specific CDR3*α* sequences, yielding sequence diversity and similarity metrics equivalent to CDR3*β* generations (**Extended Data Fig. 3a**). When evaluating the overall average similarity of all generated CDR3*β* sequences to native TCRs, TCRDiff still maintained a significantly higher sequence identity of 63.2% and an enhanced BLOSUM score of 3.52 (**Extended Data Fig. 3b**).

We then selected eight peptides whose known binding TCRs in the IMMREP23 dataset exhibit the most common CDR3*β* lengths, and we visualized their corresponding CDR3*β* sequence motifs. Across eight distinct peptide targets, TCRDiff outperformed TCR-Translate and GRATCR by generating sequence motifs with the lowest Kullback-Leibler (KL) divergences compared to the native IMMREP23 reference motifs (**Fig. 3f**). For example, TCRDiff successfully captured the canonical “R-S-S-Y” motif within the hypervariable core region of CDR3*β* loops binding to the GILGFVFTL peptide^58^ (**Fig. 3g**), as well as the characteristic “P-G-T” motif within the core region of CDR3*β* loops binding to the NLVPMVATV peptide^59^ (**Fig. 3h**). These motifs serve as functional hotspots that critically shape surface interactions with peptide residues via hydrogen bonds or electrostatic interactions (**Supplementary Fig. 4**). Furthermore, TCRDiff accurately recapitulated the binding motifs of CDR3*α* sequences, closely mirroring the original amino acid preferences (**Extended Data Fig. 3c, d**). This includes successfully capturing the conserved “X-N-N-N-D-M” motif within the core region of CDR3*α* loops recognizing the NLVPMVATV peptide^59^.

**Fig. 4.**
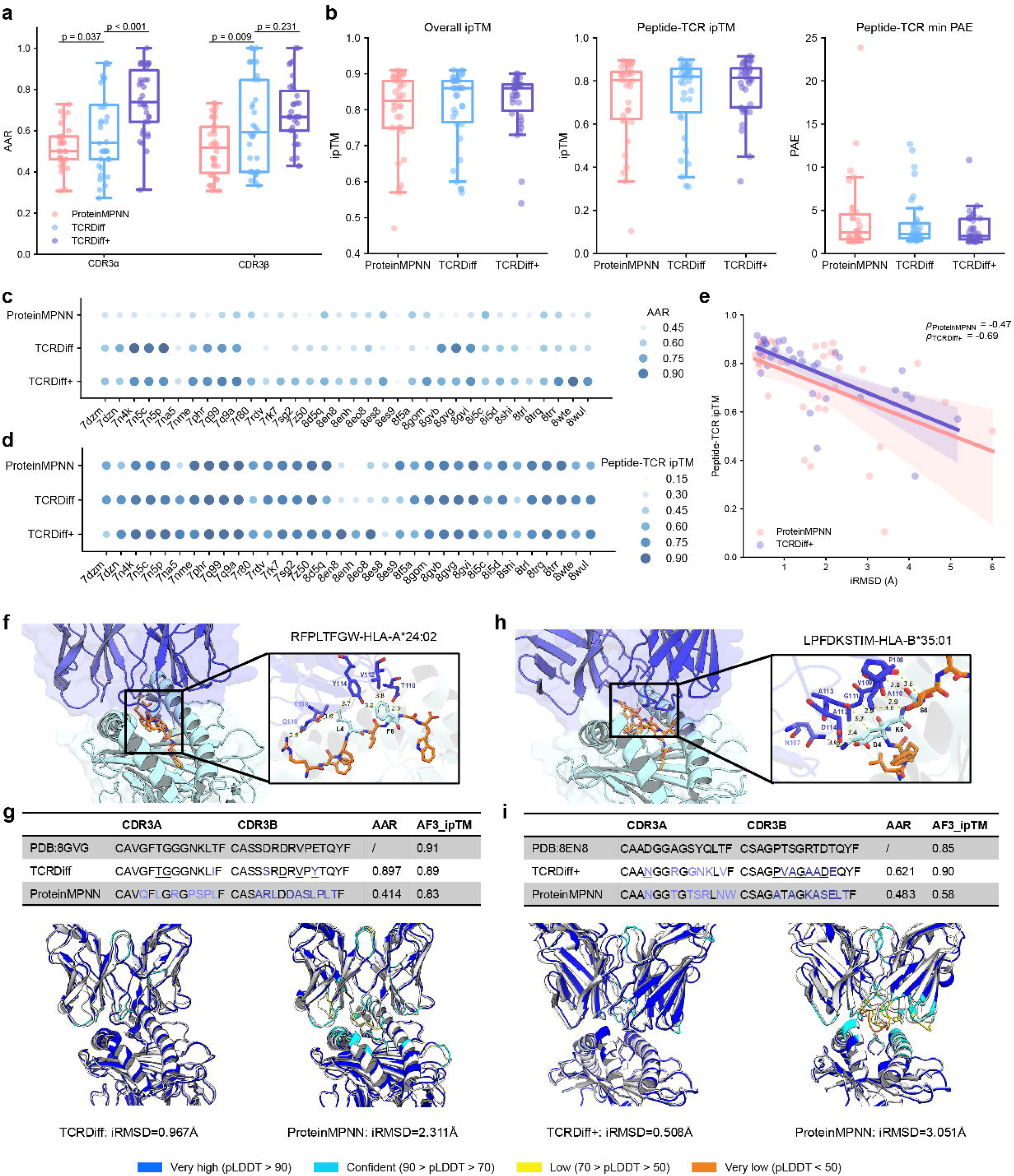
TCRDiff+ prioritizes TCR designs with enhanced sequence consistency and structural foldability. **a**, Bar plots displaying AARs for CDR3*α* and CDR3*β* loops in the STCRDab test set, generated by ProteinMPNN, standalone TCRDiff, and TCRDiff+. Statistical significance was determined using Student’s *t*-test. **b**, Bar plots evaluating structural foldability metrics (left, overall ipTM; middle, peptide-TCR ipTM; right, peptide-TCR minimum PAE) for the TCR designs on the STCRDab test set. Box center line, median; box limits, upper and lower quartiles; whiskers, 1.5×interquartile range; points, data points. **c, d**, Dot plots showing the AARs (c) and AlphaFold peptide-TCR ipTMs (d) of the TCR designs for each structural entry. Larger size and darker color of the point indicate a higher AAR or ipTM value. **e**, Correlation analysis between the peptide-TCR ipTM scores of TCR designs by TCRDiff+ and ProteinMPNN and iRMSD relative to the experimental native structures. Line, regression line; shadow, 95% confidence interval; point, data point. **f, h**, AlphaFold-predicted structures of TCR designs complexed with the RFPLTFGW-HLA-A*24:02 (f, TCRDiff) target and LPFDKSTIM-HLA-B*35:01 target (h, TCRDiff+). Insets provide zoomed-in, atomic-level visualizations of the core TCR-pMHC binding interfaces, including potential hydrogen bonds and residue contacts. **g, i**, Summary of sequence consistency and structural plausibility of TCR designs for RFPLTFGW-HLA-A*24:02 (g) and LPFDKSTIM-HLA-B*35:01 (i). AlphaFold-predicted pLDDTs of the TCR-pMHC complexes are visualized with their iRMSDs relative to the corresponding PDB structures. The underlined residues of the TCR design denote positions that form direct interactions with the bound peptide.

TCRDiff accepts flexible inputs of partial CDR3αβ sequences. When conditioning on a known sequence for one chain (CDR3*α* or CDR3*β*) and evaluating the generative performance on the other, the model achieved comparable AARs for each peptide in the IMMREP23 test set (**Supplementary Fig. 5**). To further evaluate the novelty of the generated TCR designs, we performed stochastic decoding to design TCRs targeting the six most common pMHC targets in the training set (**Supplementary Fig. 6**). When comparing these generated sequences to their most similar TCR counterparts that were excluded from the training data, TCRDiff achieved average AARs exceeding 0.8 across almost all target groups except those TCRs binding to the SARS-CoV-2 epitope TTDPSFLGRY. While the hydrophilic RAKFKQLL peptide typically restricts the diversity of CDR3*α* sequences of reactive TCRs, the TCR designs generated by TCRDiff still demonstrated robust sequence uniqueness compared to the training data.

**Fig. 5.**
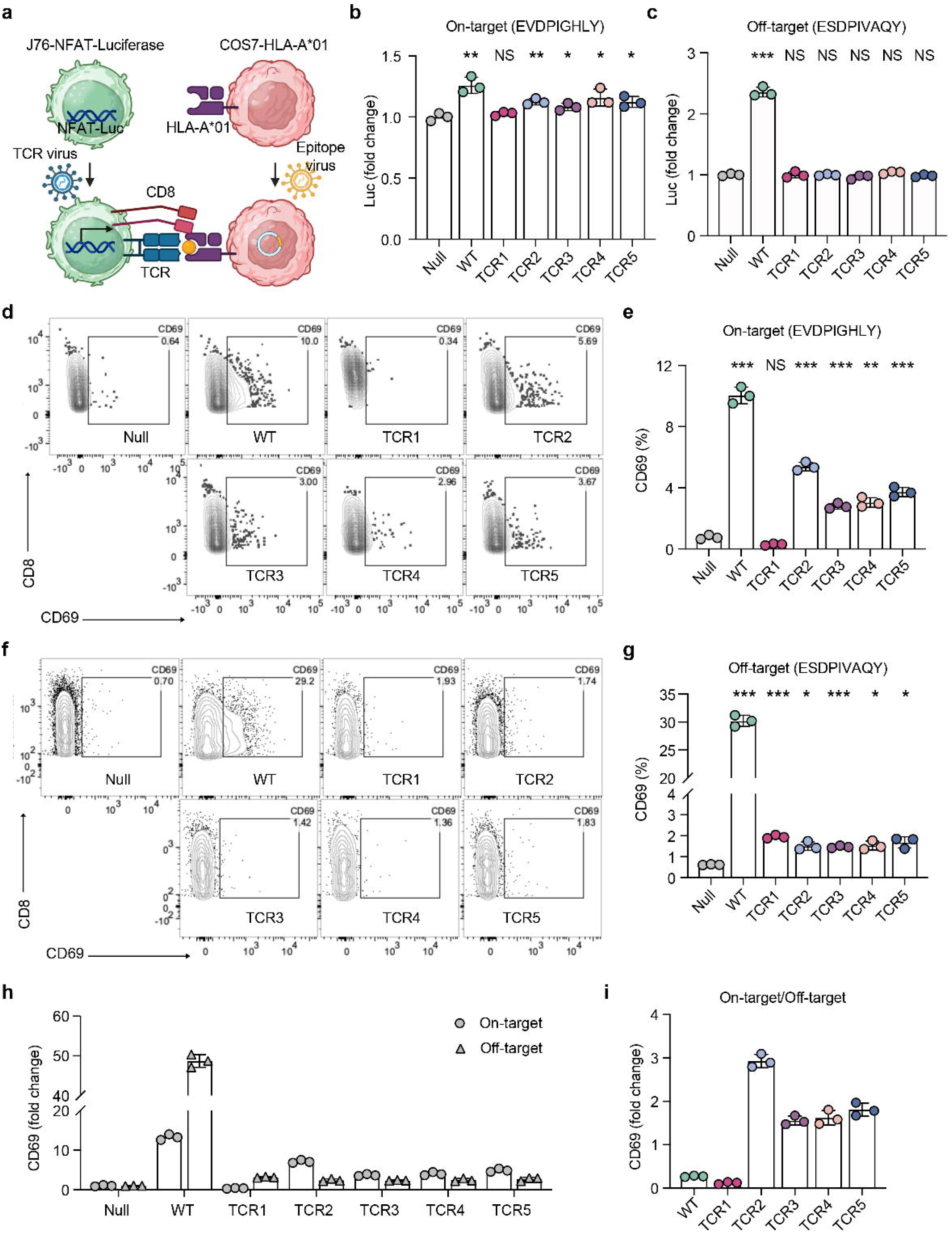
TCRDiff-designed TCRs specifically recognize the MAGE-A3 epitope in TCR-transduced reporter cells. **a**, Schematic diagram of the co-culture assay. J76-NFAT-Luciferase (plus CD8) reporter T cells transduced with the TCR virus were co-cultured with COS7-HLA-A*01 cells presenting the indicated antigen. T-cell activation was quantified by luciferase (Luc) activity fold change and flow cytometry. **b**,**c**, Fold change of luciferase activity in TCR-transduced J76 cells following co-culture with COS7-HLA-A*01 cells loaded with the on-target MAGE-A3 epitope (b) or off-target TTN epitope (c) for 6 hours. Null: no TCR transduction. **d**,**e**, Representative flow cytometry profiles of CD69 activation marker on CD8^+^ TCR-transduced J76 cells from the on-target MAGE-A3 co-culture. **f**,**g**, Representative flow cytometry profiles of CD69 activation marker on CD8^+^ TCR-transduced J76 cells from the off[target co-culture. **h**, Fold change of CD69 expression compared to the non-transduced Null control across both on-target and off-target conditions. **i**, Antigen preference quantified as the ratio of on-target to off-target activation for each designed and wild-type TCR candidate. Three independent experiments were performed. Error bar, mean ± standard deviation. Student’s t-test P values: NS, not significant; *, P<0.05; **, P<0.01;***, P<0.001.

**Fig. 6.**
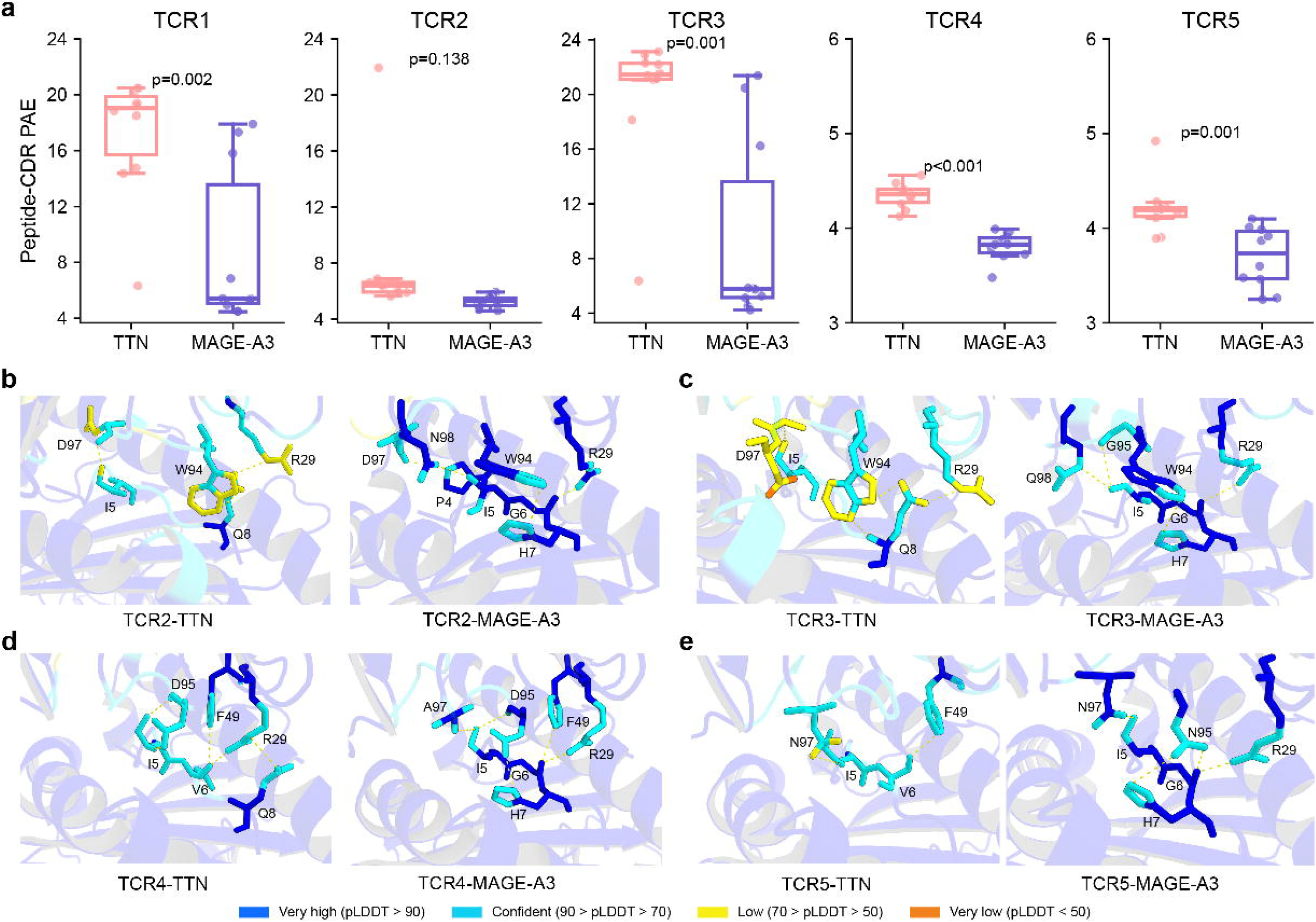
Evaluation of the structural modeling metrics and molecular interactions of the five candidate TCRs designed for specific recognition of the MAGE-A3 epitope. **a**, Average predicted align errors (PAEs) between peptide and CDR of TCR1-5 predicted by AlphaFold3 using ten random seeds. P-values were determined by Student’s t-test. Box center line, median; box limits, upper and lower quartiles; whiskers, 1.5 × interquartile range; points, data points. **b-e**, Molecular interactions within the peptide-CDR3*β* interfaces derived from the AlphaFold3-predicted TCR-pMHC complexes of TCR2 (b), TCR3 (c), TCR4 (d), and TCR5 (e). Left, off-target interface with the Titin peptide; right, on-target interface with the MAGE-A3 epitope. Dashed lines, potential residue-residue interactions within 4 Å.

### TCRDiff integrates the interface geometry to design structurally plausible TCR-pMHC complexes

We next benchmarked TCRDiff against ProteinMPNN^32^, a structure-based protein sequence design method, using the independent STCRDab test set^60^. To leverage the available 3D structural context of the TCR-pMHC binding interface, we developed TCRDiff+, which integrates the predicted logits from ProteinMPNN into the pMHC-conditioned diffusion process of TCRDiff (**Fig. 3a**). We applied a deterministic decoding strategy to generate CDR3αβ sequences and subsequently evaluated their sequence consistency compared to native TCRs and their structural plausibility via structure prediction of the designed TCR-pMHC complexes. TCRDiff significantly outperformed ProteinMPNN in the consistency of the generated CDR3αβ sequences (**Fig. 4a** and **Supplementary Fig. 7**, average CDR3*α* AAR, 0.596 vs. 0.515, P=0.037; average CDR3*β* AAR, 0.635 vs. 0.514, P=0.009). While ProteinMPNN often yielded unsatisfactory consistency with native sequences, TCRDiff+, which integrates structural knowledge from ProteinMPNN, achieved improved generative performance, particularly for CDR3*α* sequences (average CDR3*α* AAR, 0.746; average CDR3*β* AAR, 0.692). Despite relying solely on sequence inputs during model training, TCRDiff marginally outperformed ProteinMPNN in overall structural plausibility of TCR designs (**Fig. 4b**, average interface predicted Template Modeling score(ipTM), 0.805 vs. 0.795). Furthermore, the enhanced TCRDiff+ model achieved an even higher average ipTM of 0.826. The improvement in structural confidence was particularly pronounced at the critical peptide-TCR interface (peptide-TCR ipTM, ProteinMPNN: 0.716, TCRDiff+: 0.770; peptide-TCR minimum PAE, ProteinMPNN: 4.10, TCRDiff+: 2.84), where TCRDiff+ demonstrated superior metrics compared to ProteinMPNN. Taken together, these results demonstrate that TCRDiff+ achieves finer or highly comparable modeling of the peptide-TCR interface relative to standard structure-based design frameworks (**Supplementary Fig. 7**).

Across most structure entries, TCRDiff+ maintained strong sequence consistency compared to TCRDiff. Crucially, by learning from the specific docking geometry of the binding interface, TCRDiff+ effectively compensated in cases where TCRDiff alone underperformed (**Fig. 4c**). Examples include a mouse TCR in complex with HLA-A*11:01 presenting the KRAS^G12V^ peptide^61^ (PDB ID: 8WTE, AAR: 0.931 vs. 0.586), and a human SARS-CoV-2-specific private TCR in complex with RLQSLQTYV-HLA-A*02:01^62^ (PDB ID: 8GOM, AAR: 0.808 vs. 0.385). Meanwhile, the structural constraints integrated by TCRDiff+ also rescued several challenging TCR designs with initially low predicted interface confidence (**Fig. 4d**). It was evident in a cross-reactive TCR targeting an influenza NP8 epitope presented by HLA-B*35:01^63^ (PDB ID: 8EO8, peptide-TCR ipTM: 0.900 vs. 0.355), as well as a mouse TCR in complex with the HSF2 melanoma neoantigen and H2-Db (PDB ID: 7NA5, peptide-TCR ipTM: 0.825 vs. 0.715). To quantify structural alignment, we calculated the interface root mean square deviation (iRMSD) between the experimental native structures and the designed complexes (**Supplementary Fig. 8**). The peptide-TCR ipTM of the TCR complexes designed by TCRDiff+ exhibited a strong negative correlation with their corresponding iRMSD values (Pearson correlation coefficient = −0.69), validating this metric as a reliable indicator of TCR-pMHC docking quality (**Fig. 4e**). In contrast, the peptide-TCR ipTM scores derived from both TCRDiff and TCRDiff+ showed only weak positive correlations with the respective sequence AARs (**Extended Data Fig. 4a**, Pearson correlation coefficients, 0.44 and 0.27, respectively). These findings suggest that sequence consistency with the CDR3αβ sequences of native TCRs contribute to the confidence of structural predictions.

To investigate the molecular interactions within the designed TCR-pMHC complexes, we analyzed the TCR designs targeting the HIV-1 Nef138-8 peptide RFPLTFGW presented by HLA-A*24:02^64^. For this target, TCRDiff’s design exhibited better sequence consistency and structural plausibility compared to ProteinMPNN’s design (**Fig. 4f, g**). Specifically, TCRDiff substituted the peptide-interacting residue E114 in the CDR3*β* loop with Tyrosine (Y). In the predicted complex structure, this Tyrosine forms stable hydrogen bonds with F6 and favorable van der Waals interactions with the side chain of L4 of the peptide, resulting in a lower iRMSD of 0.967Å. For the TCR designs binding to the influenza epitope LPFDKSTIM (from the 1972 H3N2 strain) presented by HLA-B*35:01^63^, TCRDiff+ dramatically enhanced the predicted foldability compared to ProteinMPNN (ipTM, 0.90 vs. 0.58), even surpassing the native TCR (**Fig. 4h, j**). In this design, seven consecutive amino acids (from P108 to D114) along the hypervariable core region of the CDR3*β* loop establish a stable interaction network with multiple peptide residues. This spatial arrangement directly contributes to the remarkable improvements in both the AlphaFold pLDDT scores of the CDR3 loops and the overall iRMSD (0.508Å vs. 3.051Å).

Additionally, TCRDiff+ successfully identified key binding hotspots within the CDR3αβ loops of mouse TCRs and MHC class II-presented peptides, yielding elevated sequence consistency and highly comparable structural similarity compared to ProteinMPNN (**Supplementary Fig. 9**). Crucially, TCRDiff+ effectively rescued poor generations where standalone TCRDiff underperformed ProteinMPNN. For instance, in designing binding TCRs for the viral antigen RLQSLQTYV-HLA-A*02:01 and the neoantigen VVGAVGVGK-HLA-A*11:01 (**Extended Data Fig. 4b, c**), TCRDiff+ reconstituted the majority of amino acids along the native CDR3αβ sequences and successfully recapitulated the stable conformations within the TCR-pMHC binding interfaces.

### TCRDiff designs highly specific CDR3 sequences to recognize the MAGE-A3 epitope

To explore whether TCRDiff can generate antigen-specific TCRs with therapeutic potential, we established a TCR design pipeline that combined pMHC-conditioned CDR3αβ sequence generation, antigen binding prediction, structure-based assessment, and subsequent *in vitro* functional validation. As reducing off-target toxicity is as critical to clinical efficacy as maximizing potent T-cell activation, we selected the melanoma-associated MAGE-A3 epitope (EVDPIGHLY) presented by the HLA-A*01:01 molecule as our primary pMHC target. This choice was informed by the well-studied MAG-IC3-TCR, which exhibited clinically prohibitive off-target reactivity with a Titin (TTN)-derived self-antigen peptide (ESDPIVAQY)^44, 45^.

Utilizing different versions of our generative framework, specifically standalone TCRDiff, TCRDiff+, and a beta-chain-only model, we generated 3,000 CDR3αβ sequences. To prioritize high-specificity candidates, we applied a binding filter based on dual on-target versus off-target predicted rank percentiles, from which 19 highly confident TCR designs emerged (**Supplementary Fig. 10**). Strikingly, the majority of these generated CDR3αβ sequences displayed pronounced sequence divergence from the wild-type TCR, characterized by a Levenshtein distance of ≥ 5 amino acids (**Supplementary Fig. 11**). Following a comprehensive evaluation of structural plausibility based on AF-predicted confidence across both the predicted on-target and off-target TCR-pMHC complexes (**Supplementary Fig. 10**), we prioritized five representative TCR designs for *in vitro* experimental validation (**Extended Data Table 1**).

To characterize the TCR-mediated immune responses, the five candidate TCR designs (i.e., TCR1-5) and the wild-type MAG-IC3-TCR were transduced into J76-NFAT-hCD8A-Luciferase reporter T cell lines (**Extended Data Fig. 5a**). Anti-human CD3ε/CD28 stimulation was employed as a positive control to confirm the integrity of these engineered reporter cells (**Extended Data Fig. 5b**). These cells were then co-cultured with COS7-HLA-A*01 cells presenting either the MAGE-A3 epitope (on-target) or the TTN peptide (off-target) (**Fig. 5a, Extended Data Fig. 5c**). A Luciferase reporter assay demonstrated robust, MAGE-A3-antigen-specific activation by four out of the five designed TCRs (TCR2-5, **Fig. 5b, c**). Cells transduced with TCR2-5 exhibited a distinct increase in on-target luciferase activity (1.08-to 1.16-fold over background), while their off-target activities remained equivalent to the non-transduced negative control. In contrast, while the wild-type MAG-IC3 TCR achieved the highest on-target activity (1.26-fold), it displayed a significant off-target response (2.36-fold). Flow cytometry analysis confirmed both on-target and off-target CD69 upregulation in MAG-IC3 TCR-transduced T cells (**Fig. 5d-g**), validating its reported cross-reactivity against the human Titin protein. In contrast, TCR2-5 also exhibited substantial on-target CD69 upregulation in response to the MAGE-A3 epitope, achieving a 3.69-to 7.11-fold increase relative to the negative control (**Fig. 5h**), while maintaining marginal off-target CD69 expression. To quantify antigen preference, we calculated the relative ratio of on-target and off-target activation for all candidate TCRs. T cells transduced with TCR2-5 showed elevated CD69 upregulation upon co-culture with the cells presenting the MAGE-A3 epitope compared to those presenting the Titin peptide, yielding a 1.56-to 2.93-fold increase (**Fig. 5i**). Conversely, the wild-type MAG-IC3 TCR exhibited weaker on-target T cell activation (0.27-fold). Together, these *in vitro* functional assays validate the capacity of the TCRDiff-based generative pipeline to design and prioritize highly specific, functional TCRs with strong therapeutic potential against tumor-associated antigens.

To investigate the structural basis of this selectivity, we performed computational analyses on the predicted TCR-pMHC complex structures using AlphaFold3. All candidate TCRs exhibited superior structural plausibility when docked onto the on-target MAGE-A3 epitope presented by the HLA-A*01:01 molecule compared to the off-target Titin peptide (**Fig. 6a, Supplementary Fig. 12**). This structural advantage was localized primarily at the critical peptide-TCR/CDR interface, which dictates the antigen-binding specificity. The local geometries of the peptide-CDR3*β* interfaces revealed higher pLDDT scores and more favorable atomic contacts within the on-target complexes. For instance, Tryptophan (W94) within the CDR3*β* loop of TCR2 and TCR3 enables π-π stacking with Histidine (H7) of the MAGE-A3 epitope (**Fig. 6b, c**), enhancing the affinity and selectivity of antigen recognition^65^. Similarly, the key binding hotspots along the MAGE-A3 epitope, specifically I5-G6-H7, are specifically recognized by TCR4 and TCR5 via stronger hydrogen bonding and polar interactions (**Fig. 6d, e**). Consequently, the structural conformation of these generated TCR-pMHC complexes provides a clear geometric rationale that mirrors the observed *in vitro* T cell activation assays.

## Discussion

In this study, we have established a fundamental generative framework capable of designing highly specific, functional TCRs tailored to diverse pMHC targets. Notably, while TCRDiff possesses the unique flexibility to generate paired CDR3αβ loops across different organisms and MHC classes, it maintains competitive generative capacity compared to specialized models. The adaptive LayerNorm within the DiT blocks^46^, coupled with the PairFormer adapter^41^, enables the diffusion model to integrate conditional information smoothly. This synergistic setup effectively controls the amino acid preferences during CDR3αβ sequence generation. Conversely, conventional sequence-to-sequence (Seq2Seq) architectures rely on single-step conditioning, which often fails to capture the subtle constraints that dictate native TCR-pMHC interactions. Furthermore, by leveraging a denoising diffusion backbone, TCRDiff benefits from iterative refinement^31^ over standard autoregressive sampling, ultimately generating more structurally plausible CDR3 sequences.

As TCRDiff was fine-tuned using our pre-trained TCR-pMHC binding predictor, sequence similarities between our generated TCR designs and native binding TCRs across different MHC classes or allotypes are strongly correlated with the corresponding evaluation metrics of binding specificity. Despite a known performance skew toward antigen targets presented by highly prevalent MHC allotypes^66^, such as HLA-A*02:01, TCRDiff consistently maintained robust TCR generation quality with diverse MHC restrictions. For antigen-specific CDR3 motif deconvolution via stochastic generation, integrating explicit additional conditions, such as germline V gene usage and CDR3 loop lengths, could further enhance the fidelity of the generated motifs. This highlights the importance and value of structured conditions in the generation process.

By leveraging the structural and geometric constraints of the TCR-pMHC binding interface, we effectively integrated the ProteinMPNN logits with the TCRDiff framework, namely TCRDiff+. This integration yielded remarkable performance improvements over purely sequence-based or structure-based generative models. The improvement in sequence consistency and structural plausibility of the designed TCR-pMHC complexes might arise from model synergy, where the interface geometry guides the generation of the core regions within CDR3 loops, while the diffusion model preserves the reconstruction capacity of other conserved segments. Furthermore, AlphaFold-predicted confidence scores^47^ aligned with the structural differences between the modeled binding interfaces and their corresponding experimental structures. This alignment validates these metrics as reliable evaluation criteria for the quality of TCR designs.

While our analyses and experimental validation demonstrate that TCRDiff is a powerful computational framework for designing antigen-specific TCRs, several intrinsic limitations need to be addressed before its real-world therapeutic applications. First, the available training data for TCR-pMHC reactive pairs remains heavily biased toward prevalent viral epitopes and well-characterized tumor antigens. This data imbalance can restrict model generalizability when applied to neoantigens^49, 66^ and rare MHC allotypes, unless the model is fine-tuned on targeted experimental datasets. Furthermore, rigorously evaluating the quality of these neoantigen-specific TCR designs remains a challenge due to the scarcity of native reference TCRs for comparison. Second, since the vast majority of training TCR-pMHC pairs are derived from tetramer binding assays, candidate TCR designs optimized for the highest predicted binding affinity might not translate into successful, functional T cell activation^67^. Consequently, successful prioritization of candidate TCRs for immunotherapy still relies on a multi-tiered filtering approach that combines 3D structural modeling with *in vitro* functional validation.

Looking forward, several promising avenues can be pursued to enhance and extend the TCRDiff framework. First, incorporating refined guidance strategies into the generative diffusion process^68, 69^ could further optimize both the binding specificity and structural plausibility of the designed TCRs. For instance, external binding predictors could introduce classifier gradients during the iterative denoising diffusion steps to bias sampling toward high-affinity conformations. Alternatively, classifier-free guidance^68^ could be leveraged to control the strength of the conditioning by interpolating between antigen-specific and unconditional CDR3αβ sequence logits. Furthermore, by encoding the biophysical and geometric constraints derived from high-resolution crystal structures of TCR-pMHC complexes, structure-inspired guidance or post-generation refinement could enforce strict structural compatibility of the designed complexes with known binding modes^70^. Second, the core architecture of TCRDiff is ideally suited for the rational optimization and directed evolution of CDR loops. The framework can efficiently navigate vast mutational spaces to prioritize candidates for the experimental screening of functional paratope sites. High-throughput screening data could be iteratively fed back into the system to fine-tune the diffusion model via active learning or reinforcement learning^71, 72^. By expanding the model’s generative scope to incorporate the generation of CDR1 and CDR2 loops, TCRDiff can sever as a powerful starting point for catch-bond engineering^73, 74^, allowing researchers to explore favorable candidate mutations of the TCR interface. Finally, translating these computational designs into therapeutic applications will require systematic validation at the physiological level. A vital next step involves engineering TCRDiff-designed receptors into primary human T cells or converting them into soluble, bi-specific T-cell engagers (BiTEs). These can then undergo comprehensive *in vitro* and *in vivo* evaluations to rigorously characterize antigen-specific activation, cytokine release, tumor cell killing efficacy, and potential off-target reactivity. Through these cumulative advancements, TCRDiff will evolve into a powerful platform for rational TCR design and engineering that complements high-throughput screening experiments, ultimately accelerating the development and translation of precise TCR-based cancer immunotherapies.

## Methods

### Datasets

#### Pre-trained T cell repertoires

To establish comprehensive TCR representations, we collected diverse datasets of T cell repertoires across healthy and disease samples from humans and mice. The single-cell T cell repertoires were mainly derived from the Observed TCR Space (OTS) database^75^, 10X Genomics datasets at https://www.10xgenomics.com/datasets, a pan-disease human CD8^+^ and CD4^+^ T cell atlas (https://huarc.net/v2/atlas/)^76^, and Parse Biosciences human and mouse T cell atlas at https://www.parsebiosciences.com/resources/datasets/ (Details in **Supplementary Table 1**). After formatting V-genes and annotating germline-encoded CDR1 and CDR2 sequences of paired TCR chains, we obtained 2,475,086 distinct human TCRs and 315,078 mouse TCRs. They were then split into training, validation, and test datasets according to the ratio of 0.8:0.1:0.1 and sequence clustering of concatenated CDR3αβ sequences by MMSeq2^77^. TCRs from the healthy samples additionally made up a background dataset that was prepared for TCR binding prediction.

#### Pre-trained peptide-MHC data

We constructed an unlabeled peptide dataset based on the positive T cell and MHC ligand assays in IEDB and filtered the linear epitope sequences within the length of 8-25 amino acids, leading to 1,052,670 unique peptides. The binding affinity data between peptide and MHC class I and II molecules were derived from NetMHCpan4.1 and NetMHCIIpan 4.0^78^ (https://services.healthtech.dtu.dk/suppl/immunology/NAR_NetMHCpan_NetMHCIIpan/), respectively. The dataset is composed of 181,252 and 129,110 scaled and normalized IC_50_ values for peptide-MHC class I and II pairs, spanning 118 MHC class I allotypes and 80 class II allotypes. Protein sequences representing MHC molecules were collected from the repository of TCRdock (https://github.com/phbradley/TCRdock)^47^, and MHC pseudo-sequences of 34 amino acids were also extracted. Both the peptide sequence dataset and the peptide-MHC binding dataset were then split into training, validation, and test datasets according to the ratio of 0.8:0.1:0.1 and sequence clustering of peptide sequences by MMSeq2.

#### TCR-pMHC recognition datasets

We collected human and mouse TCR-pMHC class I and class II reactive pairs from multiple databases and studies, referring to pMTnet-omni^54^ and MixTCRpred^79^. The TCR-pMHC recognition dataset used for training was derived from IEDB^80^, VDJdb^58^, PIRD^81^, McPAS^82^, 10X^83^, and 15 studies (Details in **Supplementary Table 2**). The validation dataset was curated from another 50 studies. TCRs that contained the CDR3 sequence and V gene, at least for one chain, were retained in the datasets, and peptide targets lacking HLA restriction information were excluded. The length of CDR3 sequences was also restricted from 10 to 25 amino acids. CDR1 and CDR2 sequences were annotated based on the database of tcrdist^84^. Overlapped TCR-pMHC pairs with the validation dataset (having the same CDR3*α* or CDR3*β* sequence) were removed from the training dataset for fair evaluation. Consequently, we obtained 135,244 training TCR-pMHC reactive pairs, including 50,417 paired TRA-TRB, 64,384 TRB only, and 20,443 TRA only records, and 1,817 validation TCR*αβ*-pMHC pairs. To train and evaluate models for predicting TCR binding specificity, non-binding TCR-pMHC pairs were generated using two strategies:

1. Random shuffling within the training dataset by pairing TCRs that did not share identical CDR3*α* or CDR3*β* sequences. It produced a number of negative samples equal to three times the number of positive samples.
2. Sampling from healthy repertoires by drawing TCRs from human or mouse T cell repertoires obtained from healthy individuals. It generated a number of negative samples equal to twice the number of positive samples.

We ensured that each generated negative sample matched its corresponding positive sample in chain composition, i.e., both contained either paired TCR*α* and TCR*β* chains or a single TCR*α*/TCR*β* chain. The “non-binding” TCRs in the validation dataset were only sampled from healthy repertoires, and the sampled TCRs were then excluded in the background TCR dataset for model training. As 18 pMHCs had over 1,000 binding TCRs and made up 70.3% of the training dataset, we also constructed a subsampled dataset of 58,425 TCR-pMHC pairs in which each pMHC had a maximum of 1,000 binding TCRs. The IMMREP23 dataset^52^ from a TCR-epitope prediction challenge, which was composed of 598 positive TCR-pMHC pairs and 2,886 negative pairs covering 20 pMHC targets and 6 MHC allotypes, was employed for benchmark testing.

#### STCRDab structural test set

To evaluate the model capacity of TCRDiff and ProteinMPNN in generating CDR3αβ sequences, we established a structural test set of crystal structures of TCR-pMHC complexes curated from the STCRDab database^60^ (https://opig.stats.ox.ac.uk/webapps/stcrdab-stcrpred). We selected the complex structures deposited after 01/01/2022 and excluded the overlapped TCR-pMHC pairs within the training dataset. Finally, we obtained a structural test set of 34 TCR-pMHC complexes. The protein sequences of the complex structure and designed CDR3αβ sequences were used to predict three-dimensional structures by AlphaFold^41^.

### TCR discrete diffusion model

#### Model backbone

We employed the architecture of Diffusion Transformer^46^ (DiT) to integrate the germline-encoded V genes (represented as Atchley factors^85^ of CDR1*αβ* and CDR2*αβ* segments) into the language model to learn the biological grammar of paired CDR3αβ sequences. Specifically, adaptive LayerNorm can introduce condition-dependent scale and bias parameters in the usual LayerNorm operation, and these parameters were initialized to zero.

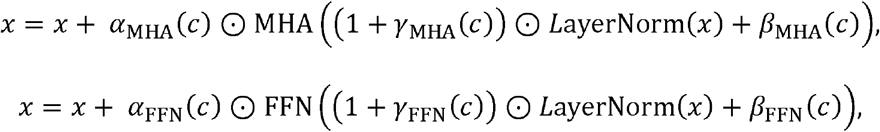

where *α,γ,β* are learnable gating, scale, and bias parameters based on the conditional embedding *c*, respectively. MHA and FFN represent the multi-head attention layer and feed-forward network in Transformer. ⊙ denotes the Hadamard product. It is compute-efficient and is restricted to applying the same function to all tokens.

We also adopted a PairFormer-like architecture inspired by AlphaFold3^41^ for modeling peptide-MHC and TCR-pMHC interactions. A PairFormer block requires single representations for protein sequences and pairwise representations that model the interaction between residue pairs. It is often composed of three major modules:

1. A Pairwise block comprising triangular multiplications, triangular gated attention layers, and a transition layer of pairwise representation. Triangle multiplication is calculated as follows:

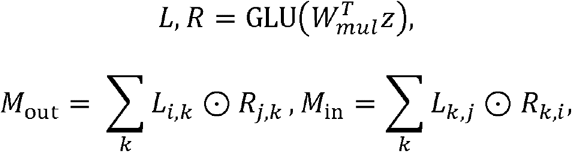

where *z* denotes the input pairwise embeddings, *M*_out_ and *M*_in_ represent multiplications according to “outgoing” and “incoming” edge orientations. After normalization, projection, and gating, the pairwise embedding are updated via residue connection. Similarly, triangular attention layers perform self-attention around the starting node and ending node of the fully-connected sequence graph:

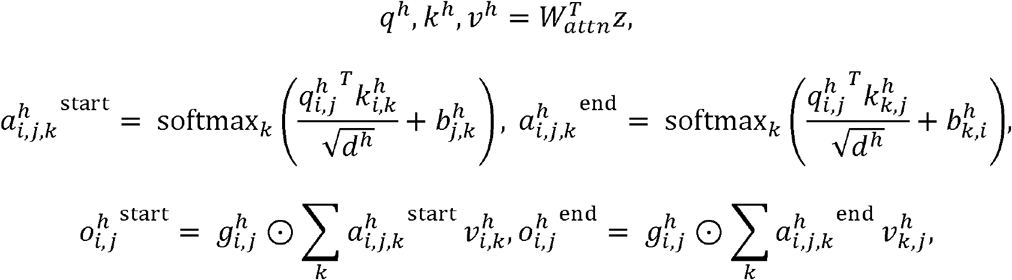

where *q*^*h*^, *k*^*h*^, *v*^*h*^ denote query, key, and value of the *h*th attention head, 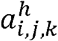 and 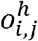 represent the attention and outputs for both attention types. 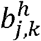 and 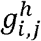 represent the learnable bias and gating parameters, respectively. *d*^*h*^ means the number of dimensions. Finally, the output of each head is concatenated, projected, and added to the pairwise embedding. The transition layer is a two-layer feed-forward network with a SwiGLU activation function^86^.
2. A self-attention layer with pairwise bias that integrates pairwise information into single representations through the attention mechanism.

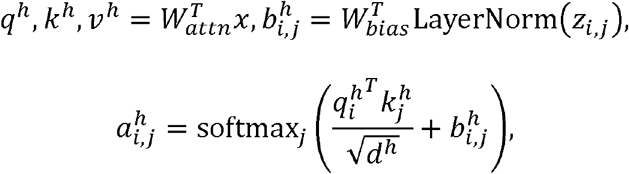

where 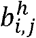 is the learnable attention bias term based on the updated pairwise embedding by the Pairwise block. Attention values are then multiplied by values and processed by concatenation, output gating, and projection. The outputs are added to the single representation.
3. The last transition layer, a two-layer feed-forward network, transforms the single representation.

The PairFormer architecture explicitly modeled interactions between residues by jointly updating single and pairwise representations and might facilitate to capture interaction patterns between TCR and pMHC.

#### Pre-trained model development

In the pre-training stage, we first developed a discrete diffusion language model for CDR3αβ sequences that incorporated conditions of germline-encoded V-genes via the DiT architecture. The forward process of discrete diffusion defines a Markov process in which the transition probabilities are as follows:

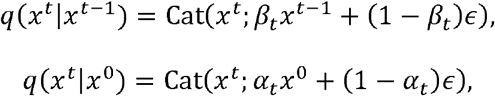

where *β*_*t*_∈[0,1) denotes the noise schedule that controls the forward corruption at timestep *t*, and 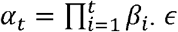 represents the stationary noise distribution. Therefore, we obtained the distribution of 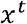 given the original data distribution *x*^0^. Like in the masked language model, we introduced the mask token in the distribution of *x*^*t*^, and set the noise distribution *ϵ* to *ϵ*( *x*^*t*^) = 1 if *x*^*t*^ ∈ *m*(*x*^*t*^) else 0 i where m(*x*^*t*^) represents the masked tokens. In this way, the forward diffusion process led to *x*^*t*^ with a masking ratio of 1 − *α*_*t*_ for each position at timestep *t*. After tokenizing the CDR3αβ sequence, the masked embedding was fed to six Transformer layers with adaptive LayerNorm. A language model (LM) head consisting of a multi-layer perceptron (MLP) and softmax activation function was utilized to predict the logits for each masked position.

The loss function of the discrete diffusion language model can simplify to the KL divergence of the true reverse distribution *q*(*x*^*t*−1^|*x*^*t*^, *x*^0^) and model reverse distribution *p*_*θ*_ (*x*^*t*−1^|*x*^*t*^) at timestep *t*:

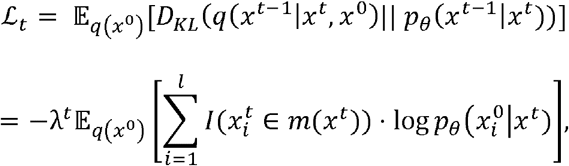

where *λ*^*t*^ is a weighting coefficient and *I*(·) is the indicator function. Therefore, the diffusion loss is transformed into a re-weighted cross-entropy loss for efficient training^31^. We first set the timestep to ensure 15% of the training CDR3αβ sequences are masked, and the model was trained with a masked language model (MLM) objective by an AdamW optimizer for 50 epochs with warm starting. In the second training stage, we randomly sampled the diffusion timesteps according to a uniform distribution to optimize the diffusion loss for another 50 epochs. We also dropped either the TCR*α* or TCR*β* chain of 30% of the TCRs in the pretrained repertoires to simulate the possible absence of one TCR chain in TCR sequencing data.

We also established a pre-trained peptide language model trained on unique peptides from the IEDB database^80^. The peptide sequences were randomly masked at a ratio of 15% and then fed into an embedding layer. A learnable positional encoding layer was used to add position information. Six Transformer layers comprising a skip-connected multi-head attention layer and a feed-forward network with standard LayerNorm were employed to capture the peptide representations. An LM head outputs the predicted logits for each masked position. Model parameters were optimized by an MLM loss for 200 epochs.

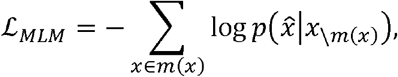

where *m*(*x*) and \*m*(*x*) denote the masked and unmasked indices of the peptide sequence.

We leveraged the pre-trained peptide language model to train a peptide-MHC binding model. The selected MHC sequences only contained membrane-distal domains (α1, α2 for class I and α1, β1 for class II) and were padded to a maximum length of 175 amino acids. MHC sequences were first transformed into BLOSUM50 encoding matrices and processed by a 1D convolutional layer. MHC representations were learned by a convolutional neural network (CNN) encoder comprising three residual 1D convolutional blocks. We extracted the MHC embeddings of the corresponding pseudo-sequences of 34 amino acids and integrated them with peptide representations via the PairFormer architecture. The pairwise representation of the concatenated pMHC sequence was initialized by dot-product attention and linear projection. Peptide and MHC embeddings were extracted from the PairFormer output. Attention pooling was performed on the MHC embedding to underscore the peptide-interacting residues, and the pooled MHC representation was concatenated to the classification embedding of the peptide. An MLP and sigmoid activation function were used to predict peptide-MHC binding affinity. In the training process, the peptide language model was also fine-tuned, and an MLM loss to reconstruct the masked tokens in 30% of peptide inputs was added to the MSE loss for predicting binding affinity.

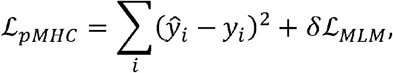

where *ŷ* and *y*_*i*_ denote the predicted and target binding affinity values, respectively. *δ* is the weighting coefficient of MLM loss.

Furthermore, we froze the parameters of the pre-trained TCR diffusion language model and peptide-MHC binding model and combined their output TCR and pMHC representations in a PairFormer adapter. To prepare the pairwise representation for residue pairs within TCR and between TCR and pMHC, dot-product attention and linear projection were performed separately. The PairFormer adapter learned the intra-chain and inter-chain interaction patterns within the TCR-pMHC pair. We obtained the single representation from the PairFormer output and split it into TCR and pMHC embeddings. A multi-head self-attention layer was further applied to each modality, and the final binding head predicted the probability of TCR-pMHC binding specificity from the concatenated classification embedding of TCR and pMHC. For each reactive TCR-pMHC pair in the training dataset, five times the non-binding TCRs to the pMHC target were generated by random shuffling and background sampling, and a binary cross-entropy loss was adopted for model training.

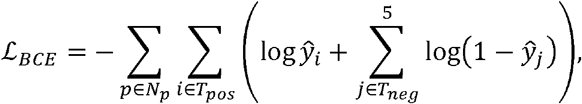

where *p* denotes a certain pMHC target, *T*_*pos*_ and *T*_*neg*_ represent the corresponding sets of binding and non-binding TCRs, respectively. *ŷ* is the predicted binding probability. The AdamW optimizer with a learning rate of 1×10^-4^ and warm starting was used to train the TCR-pMHC binding model for 80 epochs, and an early stopping strategy was employed to monitor the validation AUPR.

#### Diffusion training

Based on the architecture of the TCR-pMHC binding model, we introduced conditional information of pMHC binding preferences to the TCR diffusion language model via a PairFormer adapter, thus obtaining pMHC-conditioned TCR representations for CDR3αβ sequence decoding. The conditional diffusion training process is as follows:

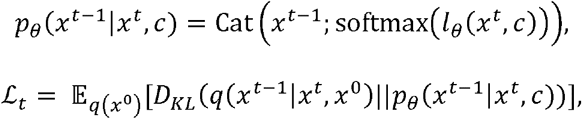

where *p*_*θ*_ (*x*^*t*−1^|*x*^*t*^, *c*) represents the conditional model and *l*_*θ*_(*x*^*t*^, *c*) is the predicted logits based on masked input *x*^*t*^ and condition *c*. The conditional diffusion language model was trained on the curated TCR-pMHC reactive pairs, and the weights of the pre-trained CDR3αβ model were fine-tuned during training to effectively adapt the model to the conditional distribution. At each training step, two coupled time steps *t*_1_ > *t*_2_ were randomly sampled from a uniform distribution. First, the ratio of *t*_1_/*T* amino acids in CDR3αβ was masked. Among these masked positions, *t*_2_ / *t*_1_ of them were unmasked at time step *t*_2_, which had less noise to ensure the consistency of the Markov process. Finally, the embeddings of 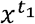 and 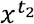 were concatenated to predict the corresponding masked amino acid. Cross-entropy loss with a linear or constant weighting strategy was utilized to train the conditional diffusion model. Model parameters were optimized by an AdamW optimizer with a learning rate of 2×10^-4^ for a maximum of 100 epochs.

#### CDR3αβ sequence decoding

TCRDiff accepts fully masked CDR3αβ with annotated V-genes and corresponding pMHC targets as the model inputs. During the generation process, we performed 10 denoising iterations along the diffusion timesteps (*t* = *T, T* − 1, … 1) following a linear or cosine schedule. At timestep *t*, the model can output a predicted categorical distribution for all amino acids along the CDR3αβ sequence.

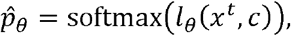

The model provides both deterministic and stochastic decoding strategies:

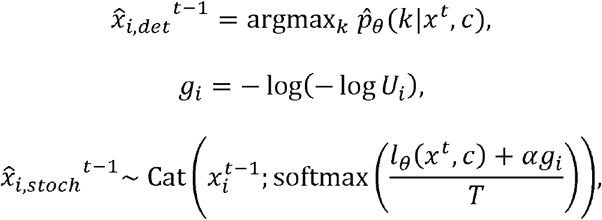

where *k* denotes all possible amino acid tokens, *U*_*i*_ represents a uniform distribution, *g*_*i*_ is the Gumbel noise, *T* is the temperature coefficient, and *α* is the noise scale. The perturbation of logits by Gumbel noise^57^ enabled the stochastic decoding of the CDR3αβ sequence. At each iteration, a certain proportion of amino acid tokens were re-masked according to the lowest *n*_1_ logit scores to simulate the noised sequence at diffusion timestep *t* − 1. The partially masked CDR3αβ sequences were then fed into the model at the next iteration, and another *n*_2_ tokens were re-masked. As *n*_2_ < *n*_1_, the model performed an iterative denoising process to retain the most confident generations at each iteration, and finally produced the completely generated sequence 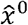. TCRDiff also supports generation tasks with partial-masked sequence inputs, including only generating the core regions in CDR3αβ or using known TCR*α*/TCR*β* chains, due to the flexible model architecture.

#### Combine ProteinMPNN logits

To optimize the binding specificity of TCR designs, the structural context of the existing TCR-pMHC binding interface can facilitate the design of functional CDR3αβ loops. We first employed ProteinMPNN^32^ to generate the CDR3αβ sequences given the structure backbone of the TCR-pMHC complex and the protein sequence of the pMHC target and the TCR germline-encoded regions. We sampled one sequence for each target in the STCRDab structural test set using a sampling temperature of 0.1. We extracted the predicted logits by ProteinMPNN and integrated them into TCRDiff logits at each iteration to provide structural guidance in the generation process.

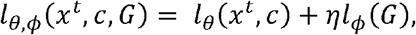

where *G* represents the input structural graph containing backbone coordinates and partial amino acid sequences used in ProteinMPNN, *l*_*θ*_ and *l* _*ϕ*_ dnote the predicted logits by TCRDiff and ProteinMPNN, respectively, and *η* is the weighting coefficient of ProteinMPNN logits to control the structural information (*η* = 0.15in our experimental setting). Deterministic and stochastic decoding strategies were both applicable to the combined logits, and we named this structure-guided TCR design approach TCRDiff+.

### Motif analysis

We explored the epitope peptides and length of CDR3αβ sequences of TCRs in the IMMREP23 dataset, and selected the most frequent combinations for motif analysis (eight for CDR3*β* and six for CDR3*α*). CDR3αβ sequences of TCRs binding to a certain pMHC target were transformed into positional information matrices. We also analyzed the generated pMHC-specific CDR3*β* sequences by TCRDiff, TCR-Translate, and GRATCR. The information matrices were visualized using the Python package logomaker^87^ to display amino acid preferences along CDR3αβ sequences. We calculated the KL divergences from the predicted motifs to those derived from the IMMREP23 test dataset to compare the generation performance.

### Structure modeling

We employed AlphaFold3^41^ to evaluate the structural plausibility of the TCR designs generated by ProteinMPNN, TCRDiff, and TCRDiff+. We kept the peptide, MHC, and germline-encoded TCR context sequences for each record in the STCRDab structural test set and replaced the CDR3αβ sequences with the generated ones. We used five random seeds to predict the three-dimensional structures for each TCR design on the AlphaFold server (https://alphafoldserver.com/), and selected the modelled structure with the highest peptide-TCR interface predicted template modelling (ipTM) in 25 AlphaFold-predicted models. The overall ipTM and minimum peptide-TCR predicted aligned error (PAE) of the predicted structure were also used to evaluate the confidence of structure prediction. We compared the structural similarity of the TCR-pMHC binding interface between predicted structures and experimental structures in the PDB database using interface root mean square deviation (iRMSD) calculated by DockQ^88^.

### TCR design pipeline

Given the wild-type MAG-IC3 TCR (PDB ID: 5BRZ), we used TCRDiff to generate 1,000 pairs of CDR3αβ sequences conditioning on the MAGE-A3 epitope, HLA-A*01:01 molecule, and TCR variable region genes (TRAV21*01, TRBV5-1*01). The lengths of the generated CDR3 sequences were the same as the length of the wild-type CDR3. Next, we employed the TCR-pMHC binding model to predict the on-target/off-target binding rank percentile of the generated TCRs against the MAGE-A3 (EVDPIGHLY) and Titin (ESDPIVAQY) peptides, respectively. In this process, 10,000 background TCRs were sampled from the T cell repertoire of healthy human samples to evaluate the relative binding potential of the generated TCRs. The binding percentile filter then yielded candidate TCR designs with high on-target binding and low off-target binding probabilities. We further employed AlphaFold to predict the three-dimensional structures of the designed TCR-pMHC complexes using 10 random seeds and calculated the confidence metrics, including TCR-pMHC ipTM, peptide-TCR ipTM, mean TCR-pMHC PAE, and mean peptide-CDR PAE. To achieve more specific binding to the MAGE-A3 peptide, we finally selected three candidate TCRs with enhanced on-target structural plausibility over the off-target ones for *in vitro* validation. We also generated MAGE-A3-specific CDR3αβ sequences using the other two strategies (by adding ProteinMPNN logits and designing CDR3*β* only) and performed similar evaluation and filtering steps to obtain another two candidate TCRs for *in vitro* validation.

### *In vitro* validation

#### Flow cytometry

J76-NFAT-Luciferase cells or COS7 cells were collected and washed once with PBS, stained with different surface antibodies in FACS buffer (PBS with 0.5% BSA) for 30□min on ice. Staining reagents include APC/Cyanine7 anti-human CD69 Antibody (FN50), APC/Cyanine7 anti-human HLA-A,B,C Antibody (W6/32), Brilliant Violet 785™ anti-human CD8 Antibody (SK1), PE/Cyanine7 anti-human TCR *α*/*β* Antibody (IP26) from BioLegend. Data were acquired on the BD LSRFortessa. All FACS data were analyzed with the FlowJo software.

#### Cell culture and cell lines

The J76-NFAT-Luciferase reporter cell line was kindly provided by Dr. Huang Huang (Stanford University School of Medicine), and the hCD8A was transduced and stored in the laboratory. COS7 cells were obtained from the ATCC and cultured under standard conditions.

#### Plasmid construction

The TCR*α* chain and *β* chain fusion gene fragment, linked by a P2A sequence, was synthesized and cloned into the pHAGE-EF1*α*-MCS-T2A-CoGFP vector. HLA-A*01 fragments were synthesized and cloned into the N103 vector (nLV dual-promoter EF-1*α*-MCS-PGK-Puro). The on-target (EVDPIGHLY) and off-target (ESDPIVAQY) epitope gene fragments were synthesized and cloned into the pHAGE-EF1*α*-MCS-IRES-BFP vector. All gene fragments were synthesized by Azenta.

#### Cell line transduction

Twelve hours before transfection, HEK-293FT cells were seeded at 2.4 × 10^6^ cells per 10-cm dish. Lentiviral supernatants were generated by co-transfecting 293FT cells with 12 μg of aim vector, 9 μg of envelope vector (psPAX2), 6 μg of packaging vector (pMD2.G), and 81 μg of PEI (YEASEN). The culture medium was replaced 6 hours post-transfection, and viral supernatants were collected 48 hours later. The supernatants were filtered through a 0.45-μm SFCA syringe filter (Corning) and concentrated by centrifugation. Concentrated virus was used to transduce J76-NFAT-hCD8A-Luciferase or COS-7 cells in the presence of 6 μg/ml polybrene (Sigma). After 48 hours, TCR expression on J76 cells and HLA expression on COS7 cells were assessed by flow cytometry, and positive cells were cultured for subsequent screening. For epitope expression, 12 μg of on-target or off-target epitope vectors were transfected into COS7 cells using 36 μg of PEI, and cells were harvested 48 hours later for detection.

#### TCR activation and luciferase assay

For TCR activation assays, 2×10^4^ COS7-HLA-A*01 cells expressing the epitope and 1×10^5^ TCR-transduced J76-NFAT-hCD8A-Luciferase cells were co-cultured for 6 hours in a 96-well plate. After co-culture, cells were harvested, and luciferase activity was measured using the Nano-Glo Luciferase Assay System (Promega). The fold induction of luciferase activity was calculated relative to unstimulated control samples. Stimulation with 5 μg/mL Ultra-LEAF™ Purified anti-human CD3 Antibody (OKT3, BioLegend) and 5 μg/mL Ultra-LEAF™ Purified anti-human CD28 Antibody (CD28.2, BioLegend) was used as a positive control.

### Statistical analysis

All statistical tests were performed two-sided in this study. Performance metrics, such as AUPR, AUC0.1, and Precision-recall curves, were computed using the Python package scikit-learn v1.7.2. The Levenshtein distances were calculated using the Python package python-Levenshtein v0.27.1. The Blosum scores of generated CDR3 sequences were calculated using the global alignment module in the Python package biopython v1.85. The sequence motifs for CDR3 sequences were plotted using the Python package logomaker v0.8.7. The ipTM and PAE matrices of predicted TCR-pMHC complexes were derived from AlphaFold prediction results, and the interface PAEs were then calculated using the Python package biopython v1.85. PyMOL v3.1.3.1 was employed to visualize the structures of TCR-pMHC complexes.

## Data availability

All data used in this study are available via Zenodo at https://doi.org/10.5281/zenodo.20586708^89^. Detailed information of the pre-trained T cell repertoires and TCR-pMHC recognition dataset is available in **Supplementary Table 1** and **Supplementary Table 2**, respectively. The IMMREP23^52^ test set was downloaded from https://github.com/justin-barton/IMMREP23/. Protein sequences and crystal structures in the STCRDab structural test set were downloaded from the STCRDab^60^ database (https://opig.stats.ox.ac.uk/webapps/stcrdab-stcrpred/Browser). The MAG-IC TCR, MAGE-A3 epitope, and Titin peptide used in the validation assays were derived from the original study^45^ and the RCSB PDB database (PDB ID: 5BRZ, 5BS0).

## Code availability

The source code and model weights of TCRDiff are available via GitHub at https://github.com/zhangyumeng1sjtu/TCRDiff and Zenodo at https://doi.org/10.5281/zenodo.20586708^89^.

### Acknowledgements

We thank Dr. Huang Huang at Stanford University School of Medicine for providing the J76-NFAT-luciferase reporter cell line. We acknowledge financial support from the National Health and Medical Research Council of Australia (Grant Nos. APP1127948, APP1144652, APP2036864 to J.S; APP2016596 to A.W.P.) and the National Natural Science Foundation of China (Grant Nos. 82541045, 82230055 to F.W.). J.S. was also supported by the Major and Seed Inter-Disciplinary Research projects awarded by Monash University.

## Author contributions

Y.Z. and J.S. conceived the ideas. Y.Z. designed and performed *in silico* experiments. W.L. designed and performed *in vitro* experiments. Y.Z., W.L., and S.X. analyzed the data and prepared figures. Y.Z. and W.L. wrote the manuscript. M.W., X.D.S., J.R., M.C.A., A.W.P., and F.W. provide guidance on data analyses. F.W. and J.S. supervised the project. All authors contributed ideas to the work and assisted in manuscript editing and revision.

## Figure legends

**Extended Data Fig. 1.**
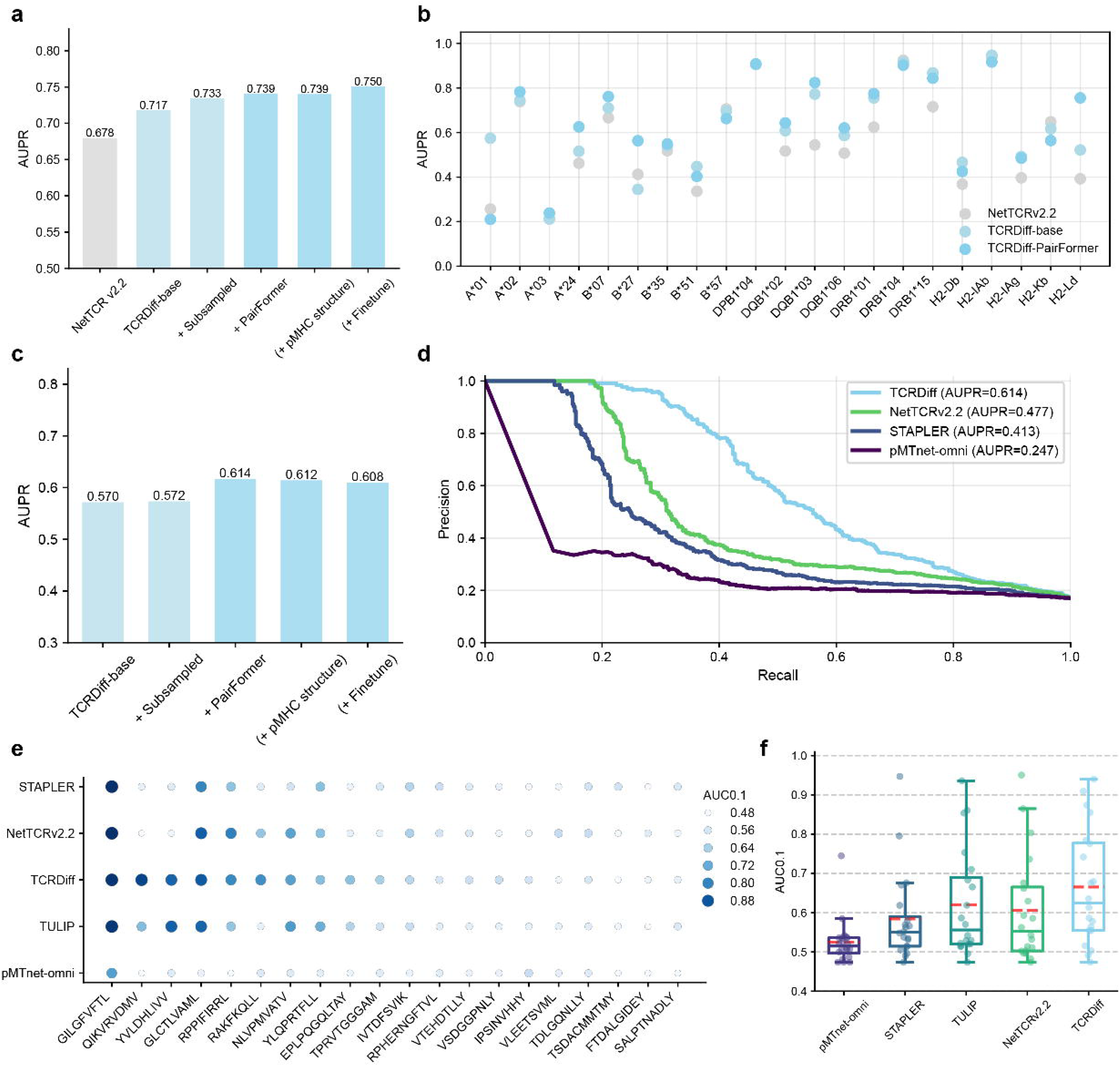
Ablation study and benchmarking on the validation set and IMMREP23 test set. **a**, Ablation study on the validation set across diverse TCRDiff model architectures and training strategies. **b**, Dot plots displaying the performance of NetTCR v2.2, TCRDiff-base, and TCRDiff-PairFormer across various validation subsets stratified by specific MHC allotypes. **c**, Ablation study on the IMMREP23 test set using TCRDiff across diverse TCRDiff model architectures and training strategies. **d**, Precision-recall curves of TCRDiff against three state-of-the-art models evaluated on the IMMREP23 test set. **e**, Dot plots displaying AUC0.1 values achieved by pMTnet-omni, STAPLER, TULIP, NetTCR v2.2, and TCRDiff across individual target peptides in the IMMREP23 test set. A larger size and darker color of the point indicate a higher AUC0.1value. **f**, Box plots displaying the distribution of AUC0.1 scores across all evaluated peptides in the IMMREP23 test set. Box center line, median; dashed line, mean; box limits, upper and lower quartiles; whiskers, 1.5 × interquartile range; points, data points.

**Extended Data Fig. 2.**
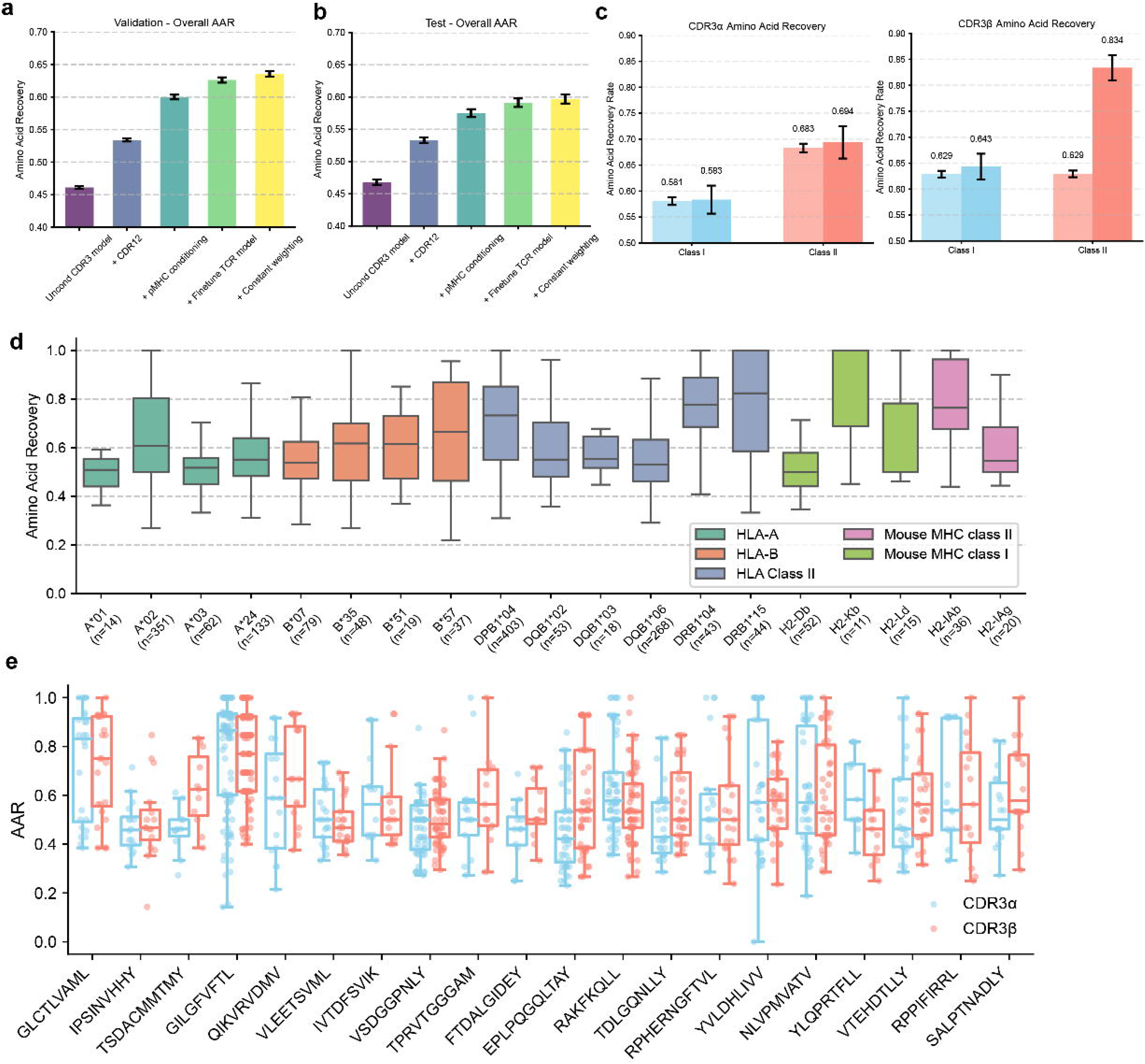
Performance evaluation of deterministic generation by TCRDiff. **a, b**, Ablation study of deterministic generation by TCRDiff using various conditional inputs and training strategies evaluated on the validation set (a) and the IMMREP23 test set (b). Bar, AAR for CDR3; error bar, standard error of AAR. **d**, Bar plots displaying the AARs of the generated TCRs compared to native CDR3*α* (left) and CDR3*β* (right) sequences, stratified by organisms and MHC classes. Error bar, standard error of AAR. **d**, Box plots displaying the distribution of AARs for TCRDiff-designed TCRs on the various validation subsets stratified by MHC allotypes. **e**, Box plots displaying the AARs of TCRDiff-generated CDR3*α* and CDR3*β* sequences across 20 distinct peptides within the IMMREP23 test set. Box center line, median; dashed line, mean; box limits, upper and lower quartiles; whiskers, 1.5 × interquartile range; points, data points.

**Extended Data Fig. 3.**
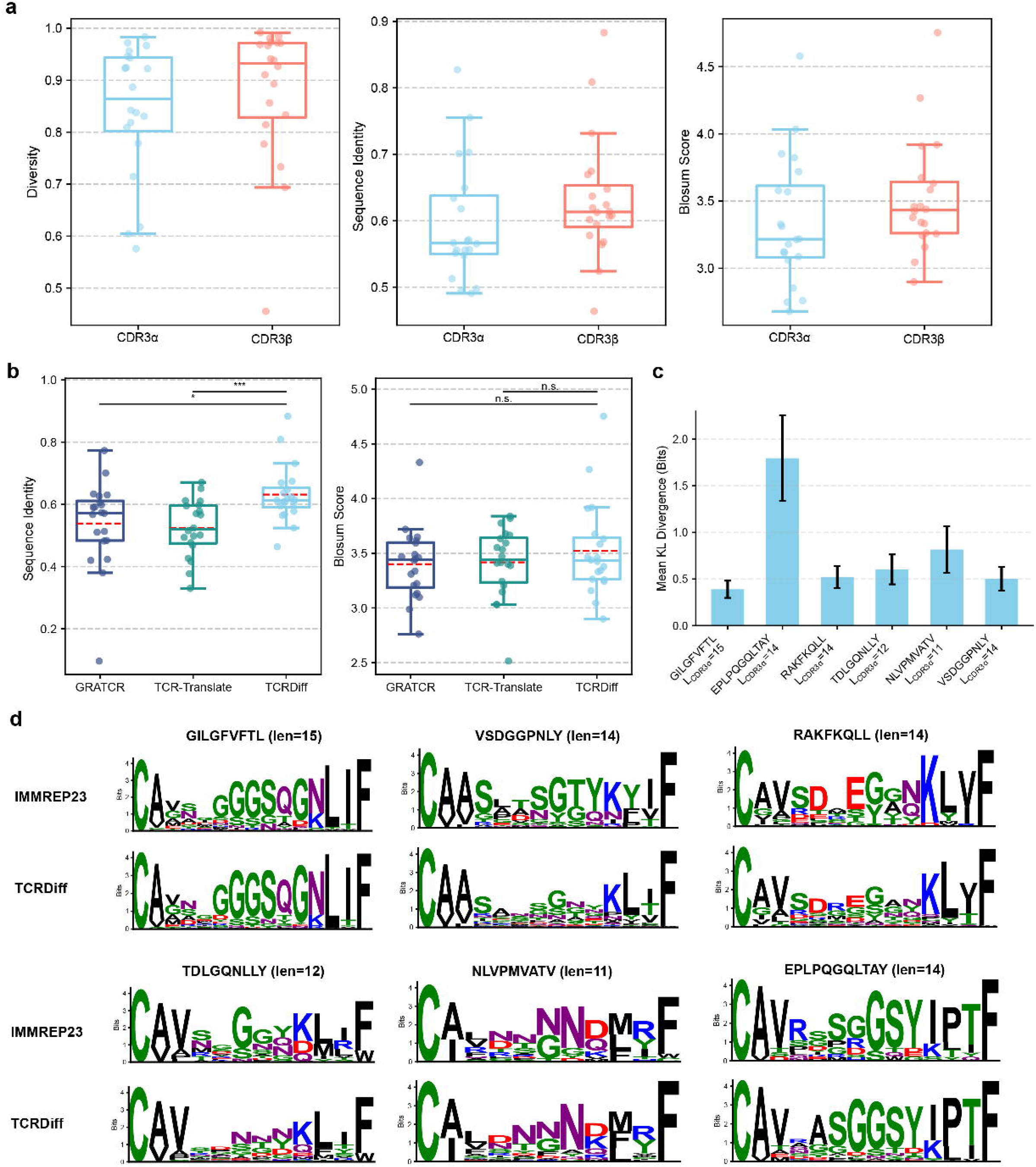
Performance evaluation and CDR3*α* motifs of stochastic generation by TCRDiff. **a**, Comparison of TCRDiff’s stochastic generation performance of CDR3*α* and CDR3*β* sequences (left, diversity; middle, sequence identity; right, BLOSUM score) on the IMMREP23 test set. **b**, Comparison of the average stochastic generation performance (left, sequence identity; right, BLOSUM score) among GRATCR, TCR-Translate, and TCRDiff on the IMMREP23 test set. Box center line, median; dashed line, mean; box limits, upper and lower quartiles; whiskers, 1.5×interquartile range; points, data points. Student’s t-test P values: *, P<0.05; **, P<0.01; ***, P<0.001; ****, P<0.0001. **c**, Mean KL divergences of TCRDiff-generated CDR3*α* motifs across various target peptides compared to the ground-truth motif derived from the IMMREP23 test set. Error bar, standard error of KL divergence. **d**, CDR3*α* motifs generated by TCRDiff and from IMMREP23 for six peptides with the most frequent length of CDR3*α*. The height of each letter is proportional to its information content.

**Extended Data Fig. 4.**
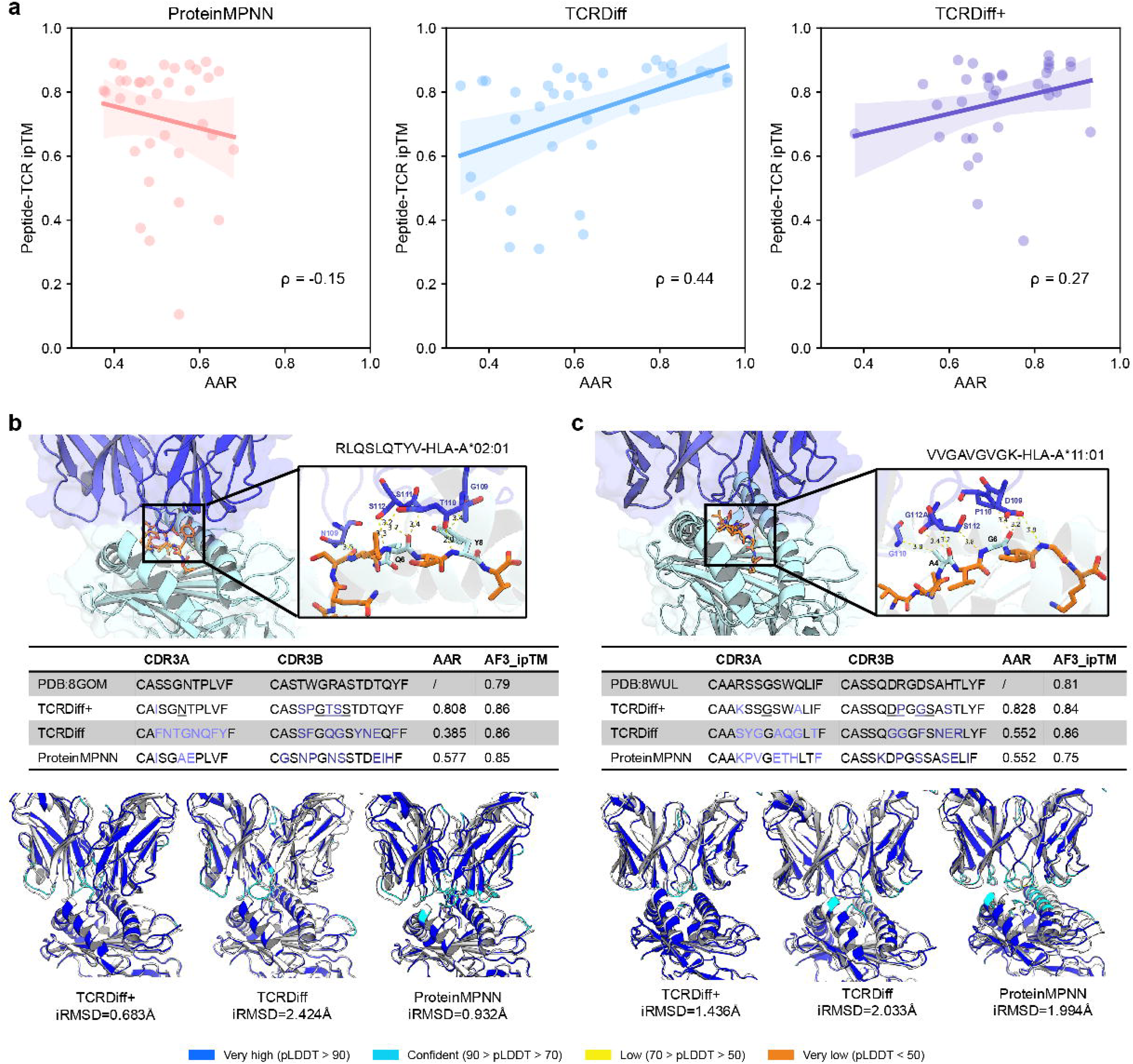
TCRDiff+ captures additional interaction hotspots ignored by TCRDiff. **a**, Correlation analysis of the relationship between peptide-TCR ipTM and AAR values for TCR designs generated by ProteinMPNN (left), TCRDiff (middle), and TCRDiff+ (right). Line, regression line; shadow, 95% confidence interval; point, data point. **b, c**, AlphaFold-predicted structures (top), summary of sequence consistency and structural plausibility of TCR designs by TCRDiff+ (bottom) binding to RLQSLQTYV-HLA-A*02:01 (b) and VVGAVGVGK-HLA-A*11:01 (c). Insets provide zoomed-in, atomic-level visualizations of the core TCR-pMHC interaction regions, highlighting potential hydrogen bonds and residue contacts. AlphaFold-predicted pLDDTs of the TCR-pMHC complexes are visualized with their iRMSDs compared to the corresponding PDB structures. Underlined residues of the TCR design denote positions that form direct interactions with the bound peptide.

**Extended Data Fig. 5.**
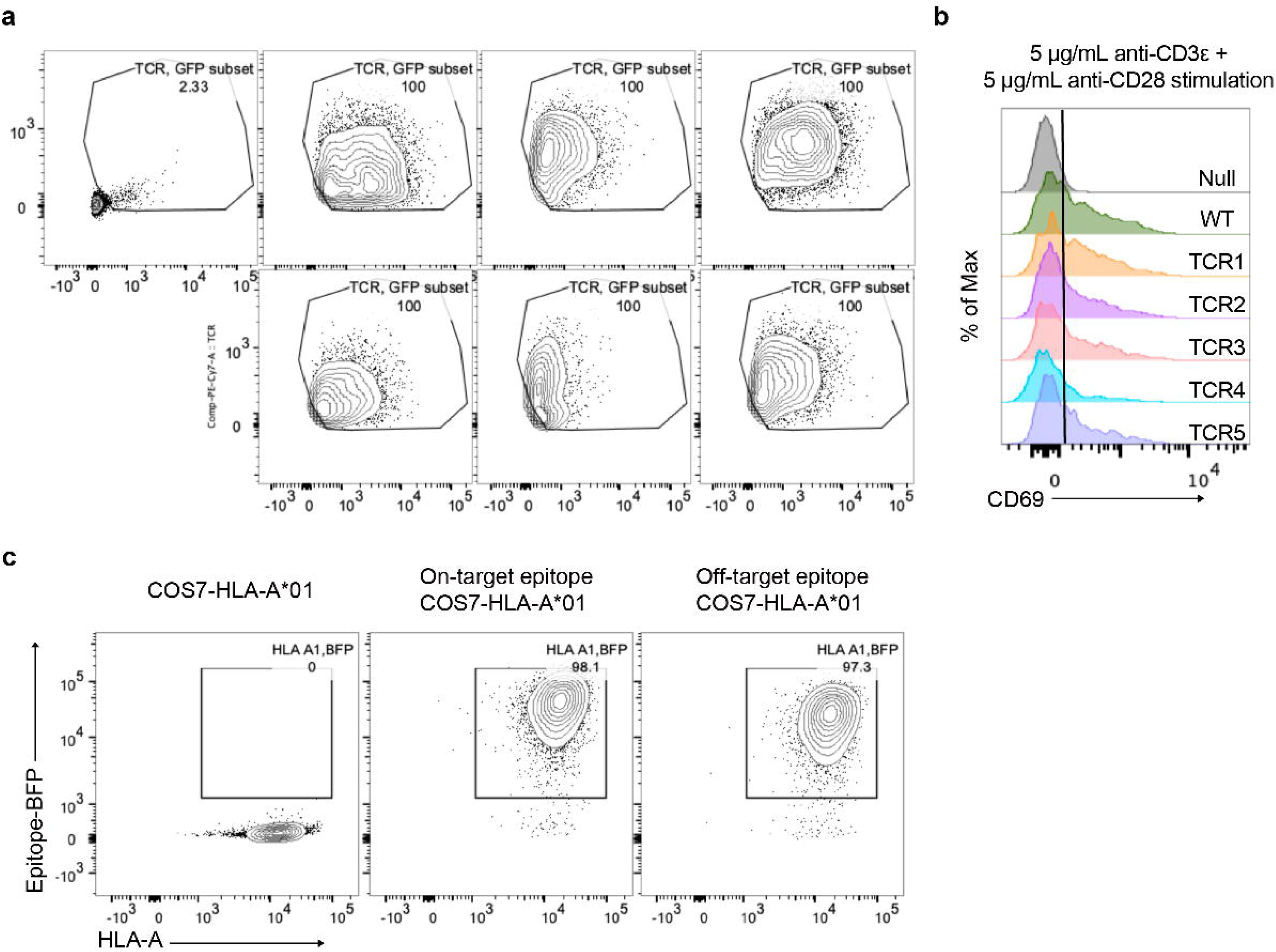
Characterization of HLA-A*01-restricted COS7 antigen-presenting cells and TCR-transduced J76-NFAT-luc reporter T cells. **a**, Flow cytometry histograms showing cell surface TCR expression (anti-TCR antibody) and GFP expression in transduced J76-NFAT-luc T cells. GFP serves as a transduction marker, while TCR surface levels indicate proper assembly and expression of the introduced TCR. **b**, Activation of TCR-transduced J76-NFAT-luc T cells after stimulation with anti-CD3ε (5□μg/mL) and anti-CD28 (5□μg/mL). CD69 upregulation was detected by flow cytometry. Histograms show CD69 expression levels in GFP□TCR^+^ gated T cells. **c**, COS7 cells expressing HLA-A*01 were loaded with either the on-target or an off-target epitope. Epitope presentation was detected via BFP fusion protein (Epitope-BFP), and HLA-A expression was measured by anti-human HLA*A,B,C antibody.

---

**Extended Data Table 1.** Summary of the on-target/off-target binding and structure prediction for the five candidate TCR designs.

| TCR | CDR3α | CDR3β | On target percentile | Off target percentile | On target Peptide-TCR ipTM | Off target Peptide-TCR ipTM | On target Peptide-CDR PAE | Off target Peptide-CDR iPAE |
| --- | --- | --- | --- | --- | --- | --- | --- | --- |
| WT | CAVRPGGAGPFFVVF | CASSFNMATGQYF | 84.89 | 67.53 | / | / | / | / |
| TCR1 | CAVRPSNFANEKLTF | CASSLDGRDGYTF | 91.90 | 31.34 | 0.825 | 0.745 | 4.504 | 6.328 |
| TCR2 | CAVPNSGTGADGLTF | CASSWGLDNEQFF | 93.15 | 48.79 | 0.840 | 0.790 | 4.604 | 5.647 |
| TCR3 | CAVRPFKAAGNKLTF | CASSWGGDQPQHF | 93.29 | 51.02 | 0.830 | 0.750 | 4.230 | 6.357 |
| TCR4* | CAVRPGGLGTDKLIF | CASSLDGADGYTF | 93.33 | 58.73 | 0.850 | 0.820 | 3.478 | 4.124 |
| TCR5** | CAVRPGGAGPFFVVF | CASSLNGNTEAFF | 96.91 | 58.37 | 0.855 | 0.825 | 3.247 | 3.891 |

Generated by \*TCRDiff+; \*\*TCRDiff (beta-only)

