## Supplementary Material for "Generative design of antigen-specific T-cell receptor sequences with a conditional diffusion model"

Yumeng Zhang^1,2,3,4,10^, Wenhua Liang^8,9,10^, Shuting Xu^1,2,3,4^, Matthew Witney^2^, Xiao-Dong Su^5^, Miles C. Andrews^6^,^7^ Jamie Rossjohn^2,3^,

Anthony W. Purcell^2,3^, Feng Wang^8,9,*^, Jiangning Song^1,2,3,4*^

**Supplementary Table 1. Data sources of pretrained T cell repertoires.**

| Data source | Organism | Data size | Raw data link |
| --- | --- | --- | --- |
| Observed TCR Space (OTS) | human, mouse | 1,680,164 human TCR, 27,154 mouse TCR | https://opig.stats.ox.ac.uk/webapps/ots/ |
| 10X Genomics datasets | human, mouse | 149,587 human TCR, 10,739 mouse TCR | https://www.10xgenomics.com/datasets |
| huARdb v2 | human | 481,550 human TCR | https://huarc.net/v2/atlas/ |
| Parse Biosciences T cell atlas | human, mouse | 384,096 human TCR, 279,033 mouse TCR | https://www.parsebiosciences.com/resources/datasets/ |

**Supplementary Table 2. Training and validation datasets of TCR-pMHC reactive pairs.**

| Data source | Train/Validation | Raw data link | Human pMHC1 pairs* | Human pMHC2 pairs | Mouse pMHC1 pairs | Mouse pMHC2 pairs |
| --- | --- | --- | --- | --- | --- | --- |
| IEDB | Train | https://www.iedb.org/database_export_v3.php | 83,985(8,068) | 3,784(1,851) | 5,577(2,293) | 557(232) |
| VDJdb | Train | https://vdjdb.cdr3.net/ | 51,061(31,611) | 2,861(1,052) | 2,907(2,318) | 505(24) |
| TetTCR-Seq | Train | https://pubmed.ncbi.nlm.nih.gov/30418433/ | 1,221(792) |  |  |  |
| TBAdb | Train | https://db.cngb.org/pird/ | 7,971(320) | 33(13) |  |  |
| TetTCR-SeqHD | Train | https://www.nature.com/articles/s41590-021-01073-2 | 4,930(4,930) |  |  |  |
| McPAS-TCR | Train | http://friedmanlab.weizmann.ac.il/McPAS-TCR/ | 9,728(1,848) | 728(26) | 98(27) | 26(0) |
| Glanville | Train | https://www.nature.com/articles/nature22976#s1 | 3,871(196) | 179(66) |  |  |
| CMV | Train | http://www.jimmunol.org/content/jimmunol/early/2018/12/23/jimmunol.1801401.full.pdf?with-ds=yes | 868(0) |  |  |  |
| Francis | Train | https://www.science.org/doi/full/10.1126/sciimmunol.abk3070 | 3,110(2,134) |  |  |  |
| Lineburg | Train | https://www.sciencedirect.com/science/article/pii/S1074761321001680 | 226(84) |  |  |  |
| Minervina | Train | https://www.nature.com/articles/s41590-022-01184-4#additional-information | 5,214(4,512) |  |  |  |
| Ishigaki | Train | https://www.nature.com/articles/s41588-022-01032-z#Sec29 |  | 83(0) |  |  |
| Tsuruta | Train | https://www.tandfonline.com/doi/full/10.1080/2162402X.2017.1415687 |  | 35(0) |  |  |
| Luo | Train | https://www.pnas.org/doi/full/10.1073/pnas.1818150116 |  | 2,146(1,231) |  |  |
| Mudd | Train | https://www.sciencedirect.com/science/article/pii/S0092867421014896 |  | 434(385) |  |  |
| Kasmani | Train | https://www.life-science-alliance.org/content/5/10/e202201503.abstract |  |  | 976(698) |  |
| 10X | Train | A new way of exploring immunity: Linking highly multiplexed antigen recognition to immune repertoire and phenotype - 10x Genomics | 17,080(17,080) |  |  |  |
| Saigusa | Train | https://www.nature.com/articles/s44161-022-00063-3 |  | 99(84) |  |  |
| Peng | Train | https://www.nature.com/articles/s41590-021-01084-z#Sec40 | 172(93) |  |  |  |
| MixTCRpred | Train | https://www.nature.com/articles/s41467-024-47461-8#Fig1 |  |  | 3,650(3,650) |  |
| NeoScreen | Validation | https://www.nature.com/articles/s41587-021-01072-6#Sec23 | 50(40) |  |  |  |
| Chen | Validation | https://www.ncbi.nlm.nih.gov/pmc/articles/PMC5472051/#SD2 | 28(28) |  |  |  |
| Orphan | Validation | https://www.ncbi.nlm.nih.gov/pubmed/29275860 | 21(21) |  |  |  |
| Ueno | Validation | http://www.jimmunol.org/content/jimmunol/169/9/4961.full.pdf | 6(6) |  |  |  |
| Grant | Validation | https://www.ncbi.nlm.nih.gov/pmc/articles/PMC5114392/ | 100(100) |  |  |  |
| Shimizu | Validation | https://www.nature.com/articles/srep03097 | 7(7) |  |  |  |
| Lichterfeld | Validation | https://academic.oup.com/intimm/article/18/7/1179/676085 | 39(11) |  |  |  |
| Motozono | Validation | https://www.ncbi.nlm.nih.gov/pmc/articles/PMC3962895/ | 19(19) |  |  |  |
| Ogunshola | Validation | https://www.nature.com/articles/s41467-018-07209-7 | 13(13) |  |  |  |
| Yu | Validation | https://journals.asm.org/doi/10.1128/JVI.01580-06 | 72(32) |  |  |  |
| p53R175H | Validation | https://www.nature.com/articles/s41467-020-16755-y | 3(3) |  |  |  |
| Peri | Validation | https://www.jci.org/articles/view/129466 | 5(5) |  |  |  |
| Sim | Validation | https://www.pnas.org/content/117/23/12826.short | 5(5) |  |  |  |
| Braunlein | Validation | https://www.ncbi.nlm.nih.gov/pmc/articles/PMC8438848/ | 7(7) |  |  |  |
| hT27 | Validation | https://onlinelibrary.wiley.com/doi/full/10.1002/eji.202049007 | 10(10) |  |  |  |
| Sooda | Validation | https://researchrepository.murdoch.edu.au/id/eprint/54110/ | 19(19) |  |  |  |
| Rowntree | Validation | https://www.jimmunol.org/content/205/6/1524.abstract | 24(23) |  |  |  |
| Wagner | Validation | https://www.sciencedirect.com/science/article/pii/S2211124721017186#app2 | 6(6) |  |  |  |
| Wu | Validation | https://www.nature.com/articles/s41467-021-27669-8#additional-information | 5(5) |  |  |  |
| Mallajosyula | Validation | https://www.science.org/doi/epdf/10.1126/sciimmunol.abg5669 | 2(2) | 2(2) |  |  |
| Nguyen | Validation | https://www.ncbi.nlm.nih.gov/pmc/articles/PMC8049468/ | 18(17) |  |  |  |
| Brunk | Validation | https://onlinelibrary.wiley.com/doi/10.1002/eji.202149290 | 13(13) |  |  |  |
| Ciacchi | Validation | https://www.sciencedirect.com/science/article/pii/S002192582200059X |  | 5(5) |  |  |
| Stadinski | Validation | https://www.nature.com/articles/s41590-019-0414-1#Sec27 |  |  |  | 24(24) |
| Watson | Validation | https://www.sciencedirect.com/science/article/pii/S2211124720308664#bib40 |  | 2(2) |  |  |
| Lu | Validation | https://rupress.org/jem/article/218/12/e20211327/212701/Identification-of-conserved-SARS-CoV-2-spike |  | 11(11) |  |  |
| Assmus | Validation | https://www.ncbi.nlm.nih.gov/pmc/articles/PMC7511415/pdf/nihms-1618744.pdf |  |  | 46(46) |  |
| Dahal | Validation | https://www.jbc.org/article/S0021-9258(20)40029-8/fulltext |  | 7(7) |  |  |
| Uchida | Validation | https://www.sciencedirect.com/science/article/pii/S2352345X20300631 |  | 23(19) |  |  |
| Schinkelshoek | Validation | https://www.sciencedirect.com/science/article/pii/S0165572818305368 |  | 20(14) |  |  |
| Luo2 | Validation | https://www.frontiersin.org/articles/10.3389/fimmu.2020.00623/full | 3(3) |  |  |  |
| Poncette | Validation | https://www.jci.org/articles/view/120391#sd |  | 10(10) |  |  |
| Wahl | Validation | https://www.science.org/doi/full/10.1126/sciimmunol.abm9644 |  | 64(64) |  |  |
| DeWitt | Validation | https://www.ncbi.nlm.nih.gov/pmc/articles/PMC6162092/pdf/elife-38358.pdf | 43(43) | 21(21) |  |  |
| Huisman | Validation | https://academic.oup.com/jid/advance-article/doi/10.1093/infdis/jiaa512/5893824?login=true | 7(7) |  |  |  |
| Cobo | Validation | https://www.sciencedirect.com/science/article/pii/S1465324922000792 | 17(17) |  |  |  |
| Moore | Validation | https://www.science.org/doi/full/10.1126/sciimmunol.abj4026 | 4(4) |  |  |  |
| Hanada | Validation | https://www.sciencedirect.com/science/article/pii/S1535610822001271 | 1(1) | 3(3) |  |  |
| Foy | Validation | https://www.nature.com/articles/s41586-022-05531-1#Sec4 | 37(37) |  |  |  |
| Axelrod | Validation | https://www.nature.com/articles/s41586-022-05432-3#Sec29 | 1(1) |  | 3(3) |  |
| Luo3 | Validation | https://www.pnas.org/doi/full/10.1073/pnas.2205797119 |  | 255(255) |  |  |
| Ma | Validation | https://pubmed.ncbi.nlm.nih.gov/36130828/ | 4(4) |  |  |  |
| Ting | Validation | https://www.pnas.org/doi/full/10.1073/pnas.1914308117 |  | 22(22) |  |  |
| TCRdock | Validation | https://github.com/phbradley/TCRdock | 117(117) | 37(37) | 39(39) | 27(27) |
| Rowntree2 | Validation | https://www.pnas.org/doi/abs/10.1073/pnas.2411428121 | 313(194) | 125(85) |  |  |
| Zhang | Validation | https://www.nature.com/articles/s41590-023-01508-y#Abs1 | 98(43) | 230(154) |  |  |
| Ciacchi2 | Validation | https://www.cell.com/immunity/fulltext/S1074-7613(23)00140-1 |  | 181(87) |  |  |
| Nguyen2 | Validation | https://www.cell.com/cell-reports-medicine/fulltext/S2666-3791(23)00127-1#mmc2 | 186(110) | 196(162) |  |  |
| Zdinak | Validation | https://www.nature.com/articles/s41592-024-02255-0 |  |  |  | 17(17) |
| Ford | Validation | https://www.nature.com/articles/s41590-023-01692-x#Fig5 | 11(11) |  |  |  |

*The values in brackets denote the numbers of complete TCRαβ-pMHC records.

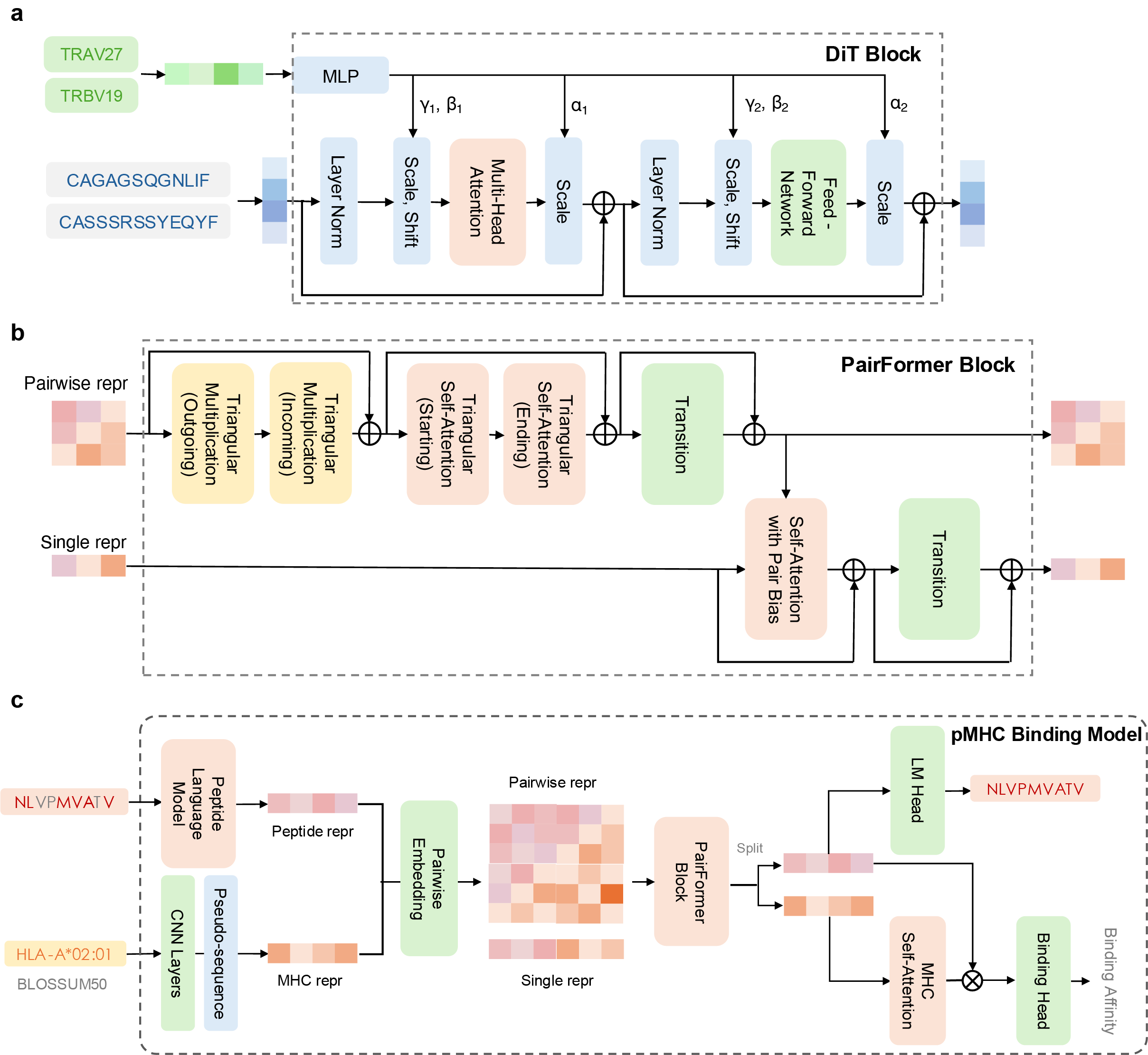

**Supplementary Fig. 1. Model architectures of the essential modules in TCRDiff.** **a,** DiT block, **b,** PairFormer block, and **c,** pMHC binding model.

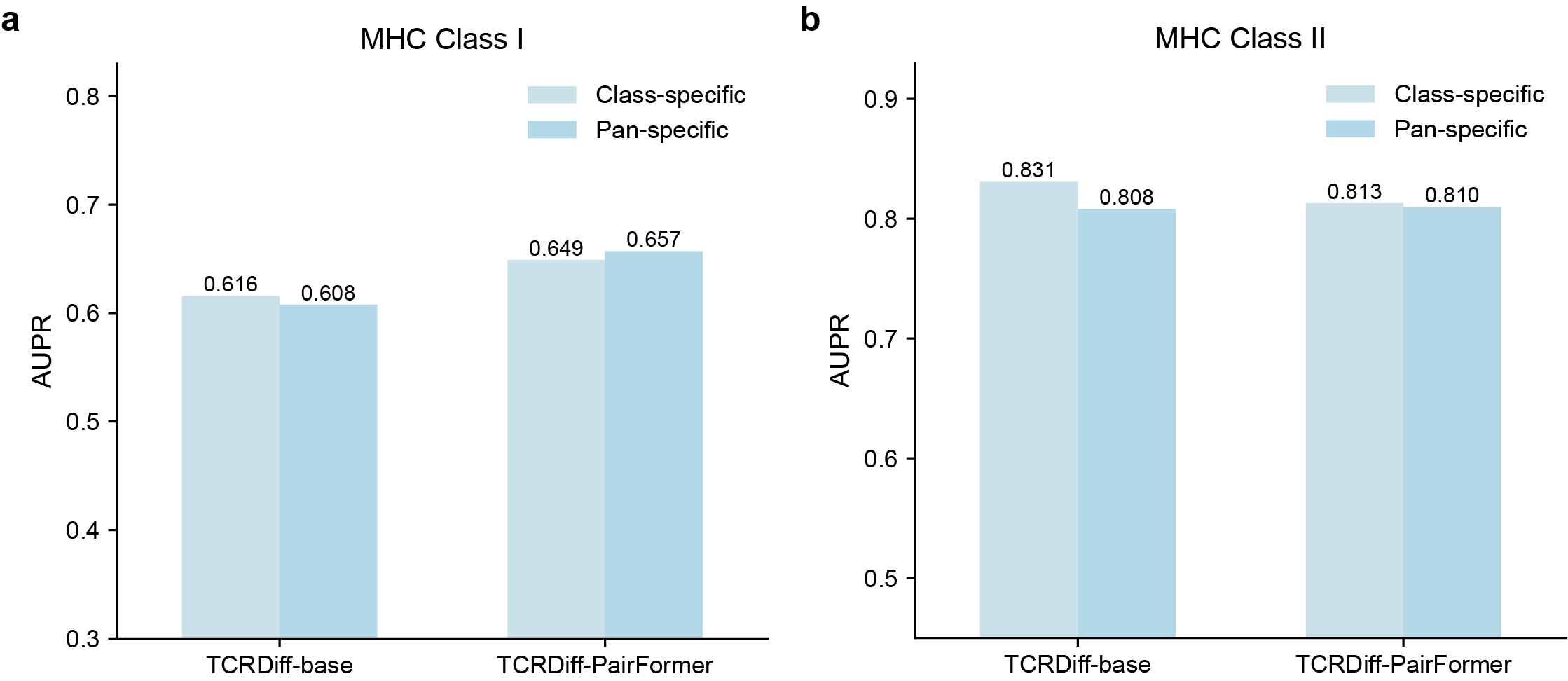

**Supplementary Fig. 2. Performance comparison on the validation subsets.** **a,** MHC class I and **b,** MHC class II between pan-specific and class-specific TCRDiff models.

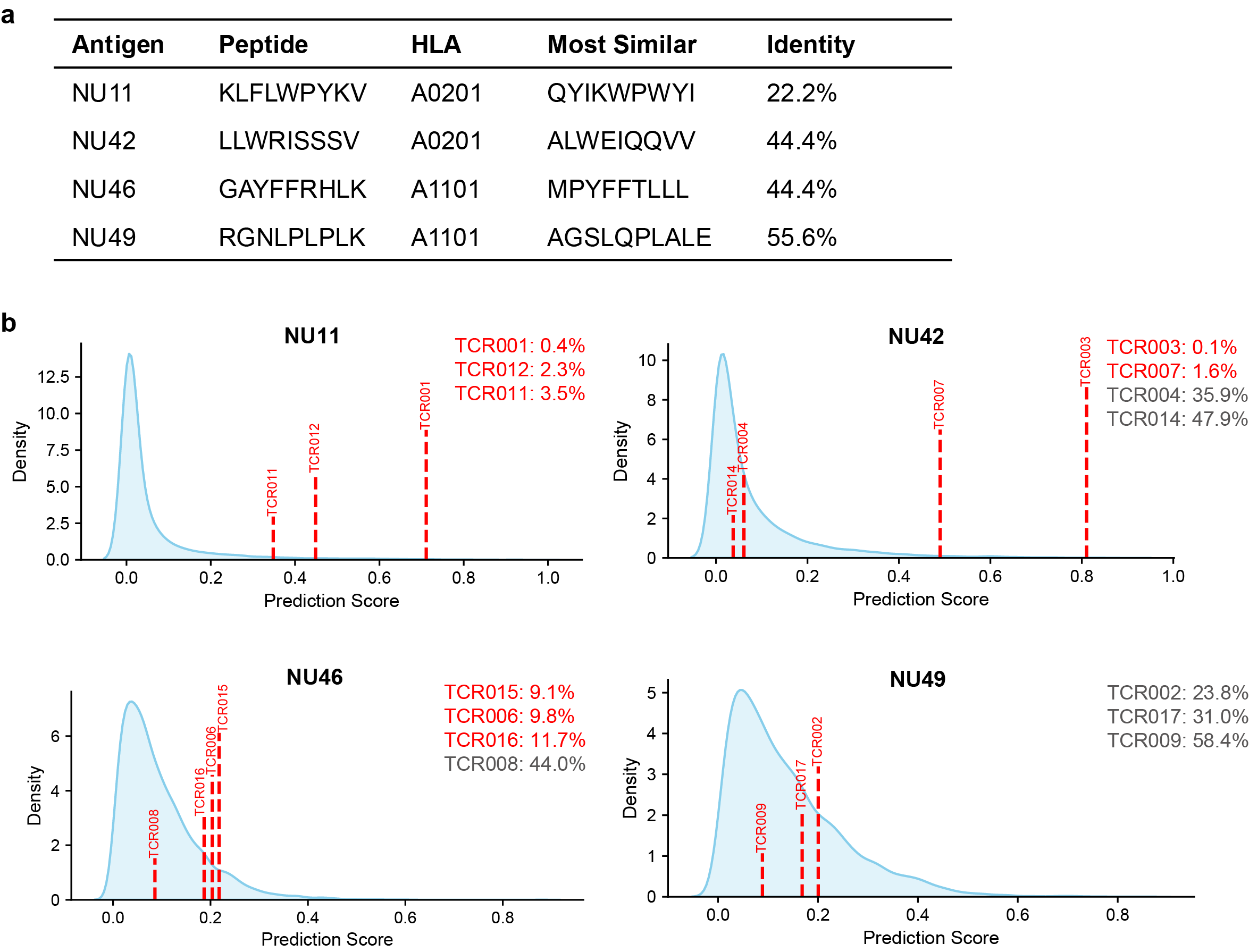

**Supplementary Fig. 3. TCRDiff identifies reactive TCRs targeting noncanonical pHLAs. a,** Statistics of four cancer-restricted cryptic antigens. **b,** TCRDiff identifies several reactive TCRs binding to these noncanonical neoantigen targets.

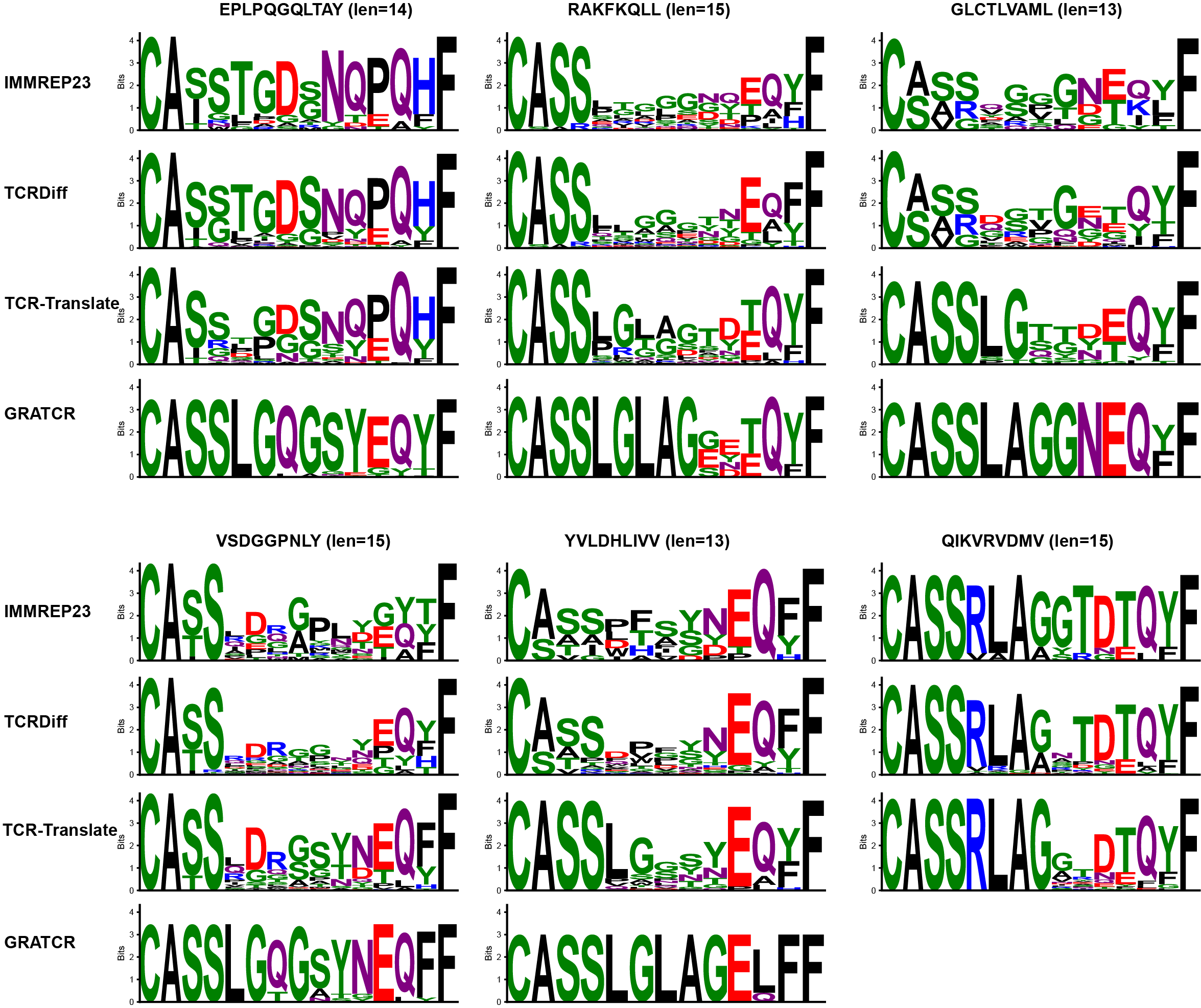

**Supplementary Fig. 4.** **Motif analysis for another six peptides in the IMMREP23 test set.** CDR3β motifs generated by the GRATCR, TCR-Translate, TCRDiff, and from IMMREP23 for another six peptides in the IMMREP23 test set with the most frequent length of CDR3β. The height of each letter is proportional to its information content.

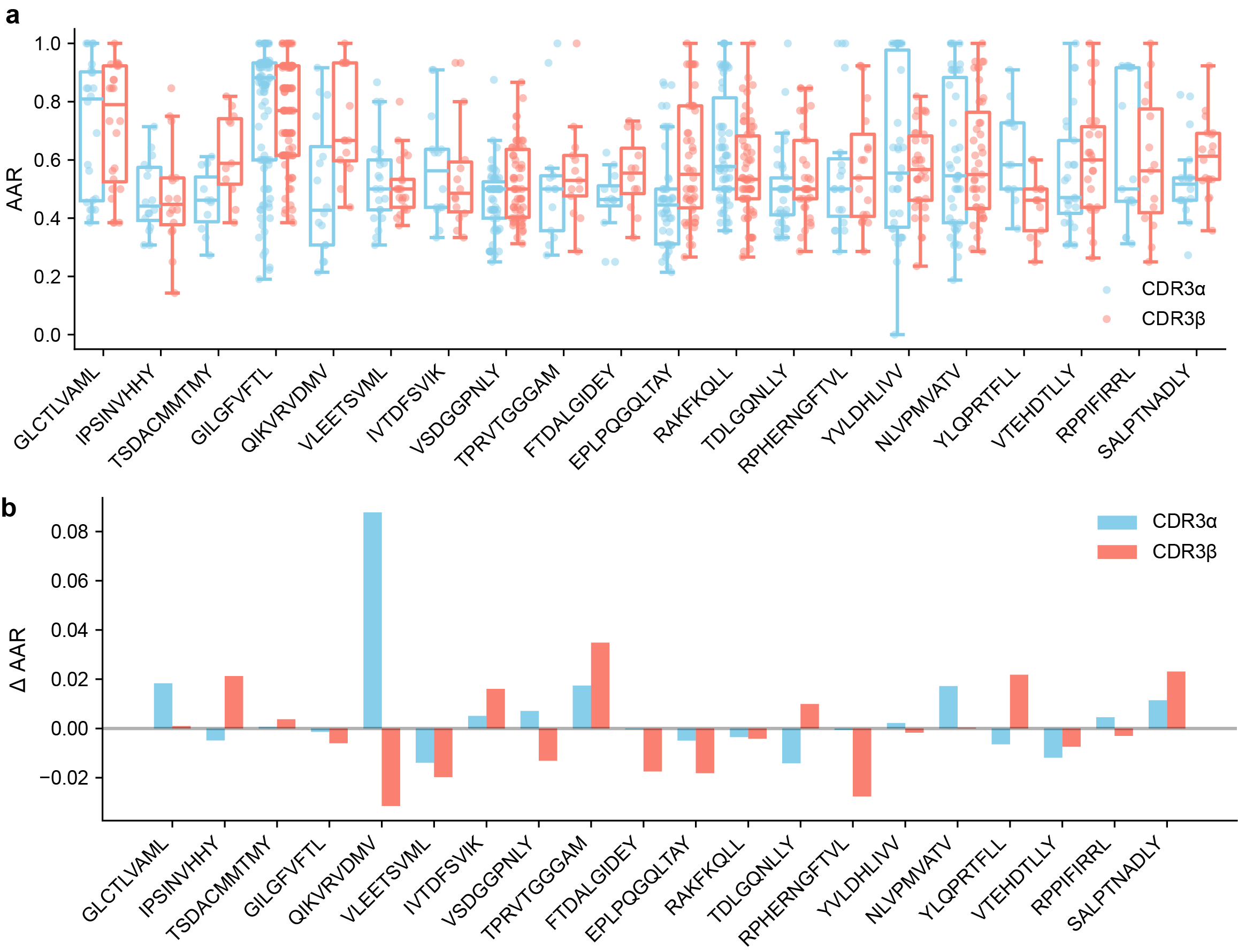

**Supplementary Fig. 5.** **Generation performance of TCRDiff given one TCRα/TCRβ chain. a,** Box plots displaying the AARs of partial chain-generated CDR3α and CDR3β sequences on 20 peptides in the IMMREP23 test set. Box center line, median; dashed line, mean; box limits, upper and lower quartiles; whiskers, 1.5 × interquartile range; points, data points. **b,** Bar plots showing the difference in AAR compared to fully-masked generations.

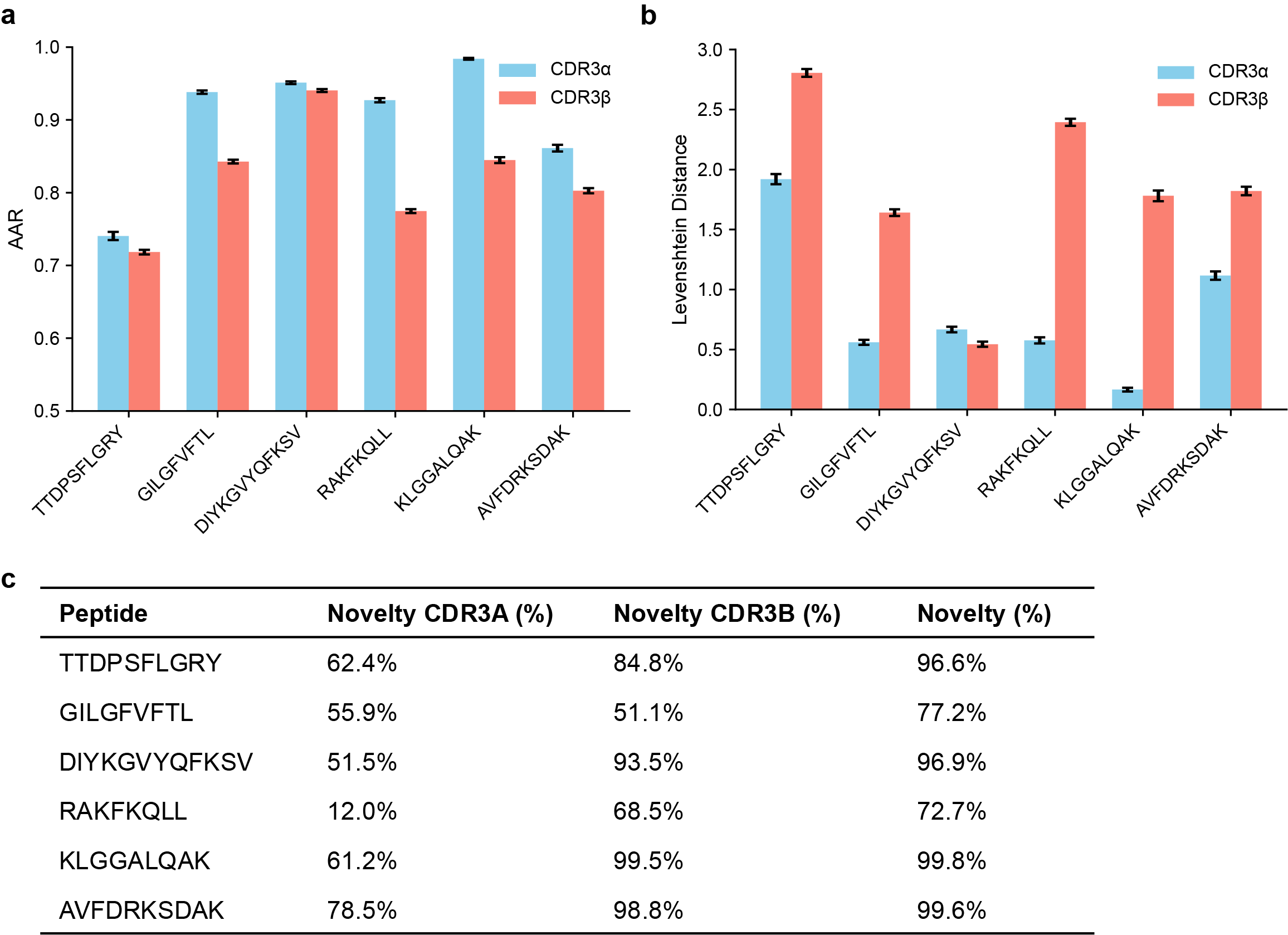

**Supplementary Fig. 6. Stochastic decoding results of TCRs binding to the most common peptides.** **a, b,** Sequence consistency (a) AAR and (b) Levenshtein distance of TCRDiff-generated TCRs for the most common peptide targets. Bar, average value; error bar, standard error. **c,** Novelty of the generated CDR3α and CDR3β sequences compared to training data.

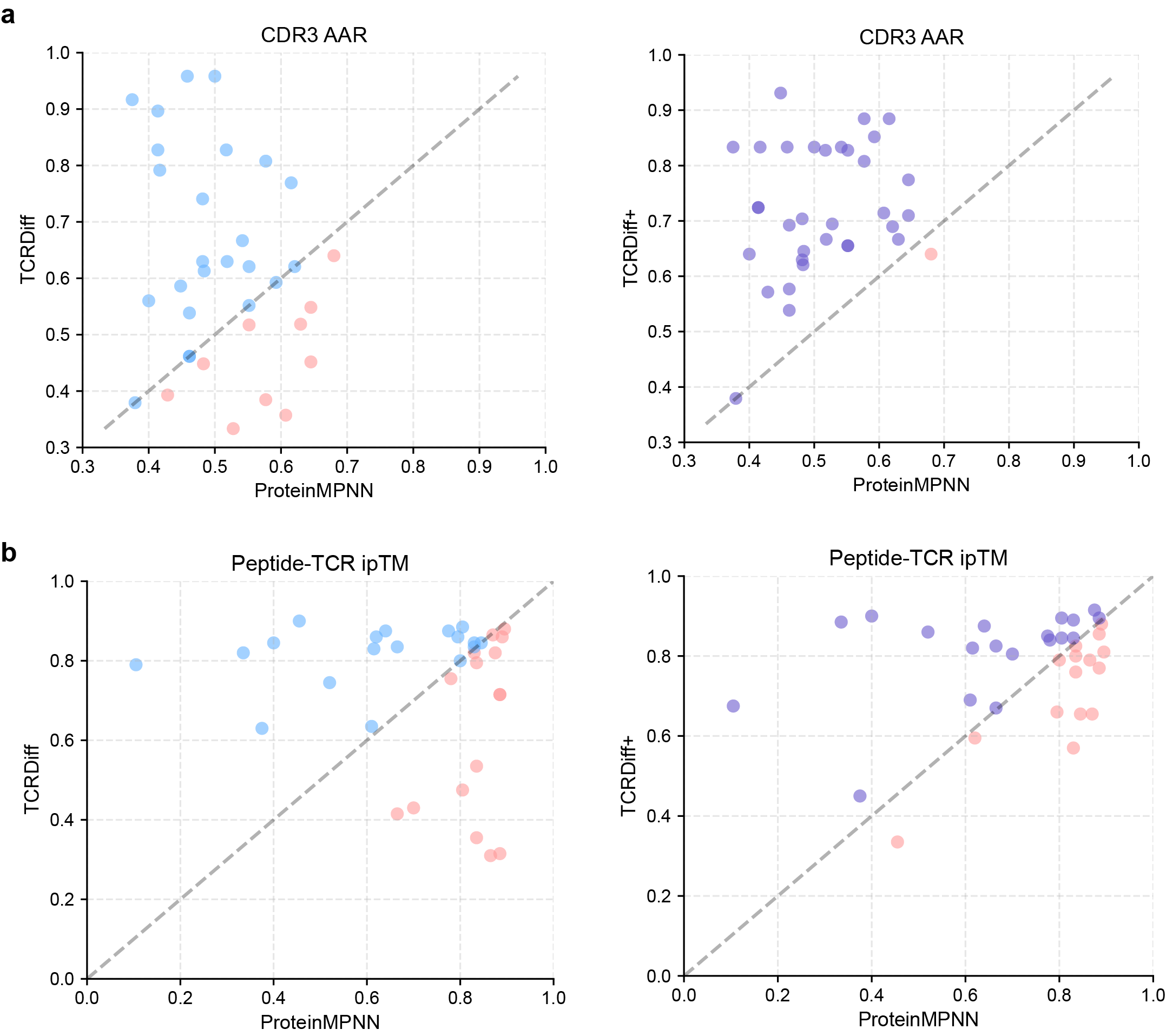

**Supplementary Fig. 7.** **Pairwise comparison of generation quality. a,** AAR and **b,** peptide-TCR ipTM. Pink, blue, and purple points indicate that ProteinMPNN, TCRDiff, and TCRDiff+ achieve better performance, respectively.

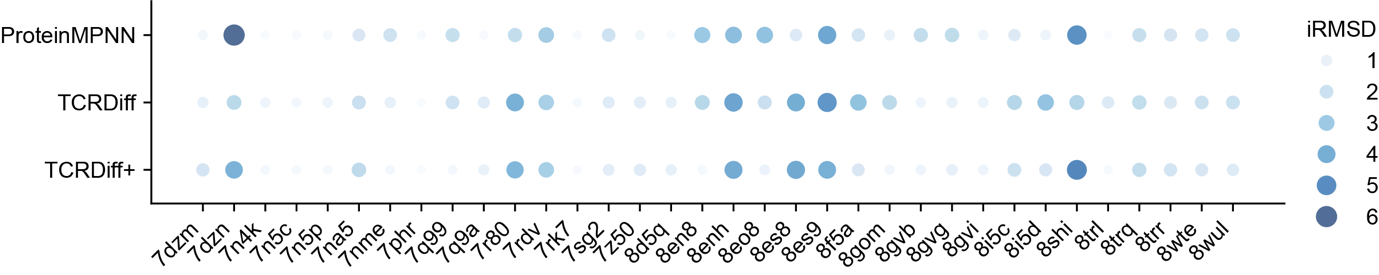

**Supplementary Fig. 8. Similarity between experimental and designed structures.** Dot plots showing interface RMSD of the TCR designs by ProteinMPNN, TCRDiff, and TCRDiff+ for each structural entry. Larger size and darker color of the point indicate a higher iRMSD value.

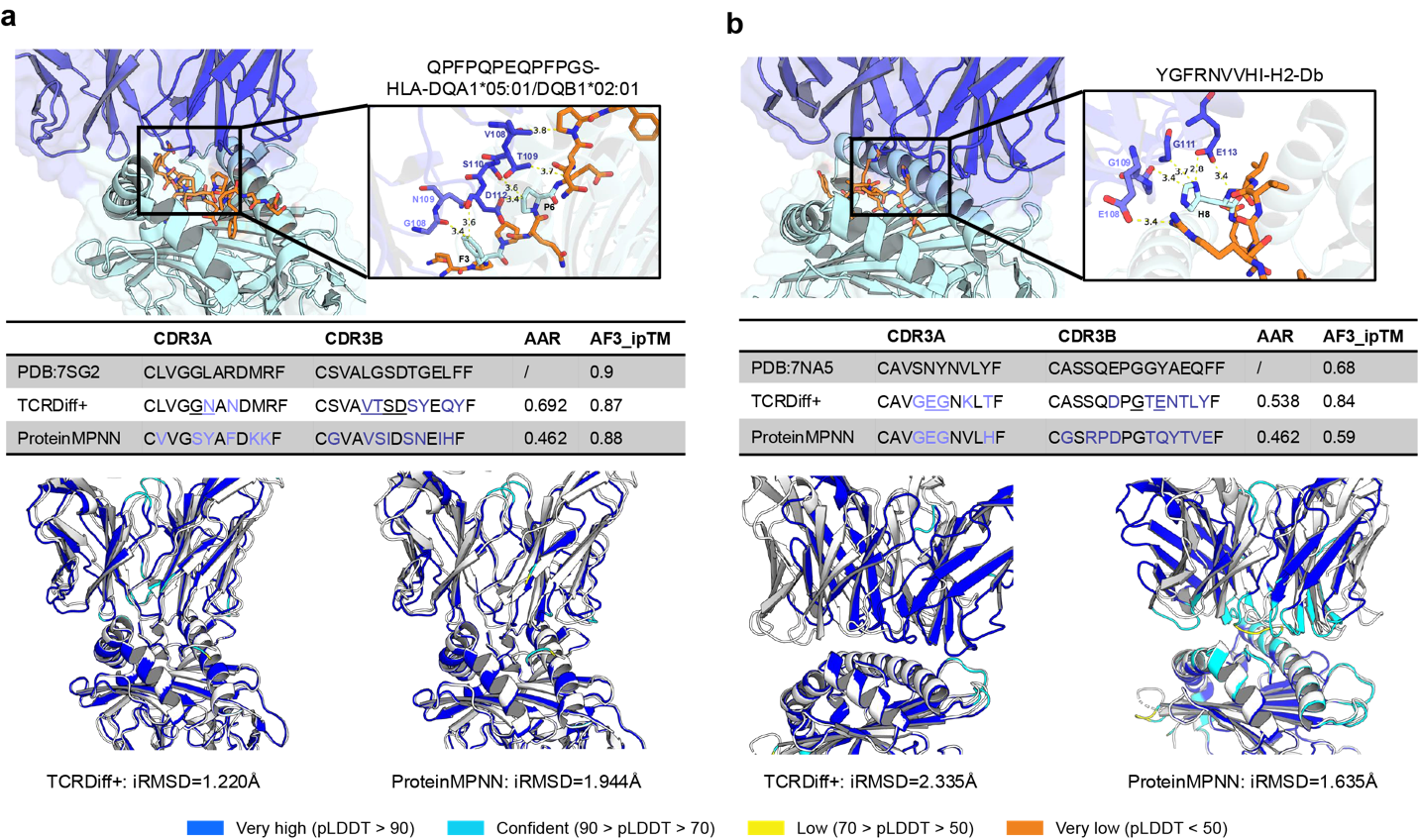

**Supplementary Fig. 9. TCRDiff+ generalizes to design TCRs for MHC class II-presented peptides or Mouse MHC.** AlphaFold-predicted structures (top), summary of consistency and structural plausibility of TCR designs by TCRDiff+ (bottom) binding to QPFPQPEQPFPGS-HLA-DQA1*05:01/DQB1*02:01 (a) and VGFRNVVHI-H2-Db (b). The core TCR-pMHC interaction regions, including potential hydrogen bonds and residue contacts, are visualized by zoomed-in views. AlphaFold-predicted pLDDTs of the TCR-pMHC complexes are visualized with their iRMSDs compared to the corresponding PDB structures. The underlined residues of the TCR design form interactions with the peptide.

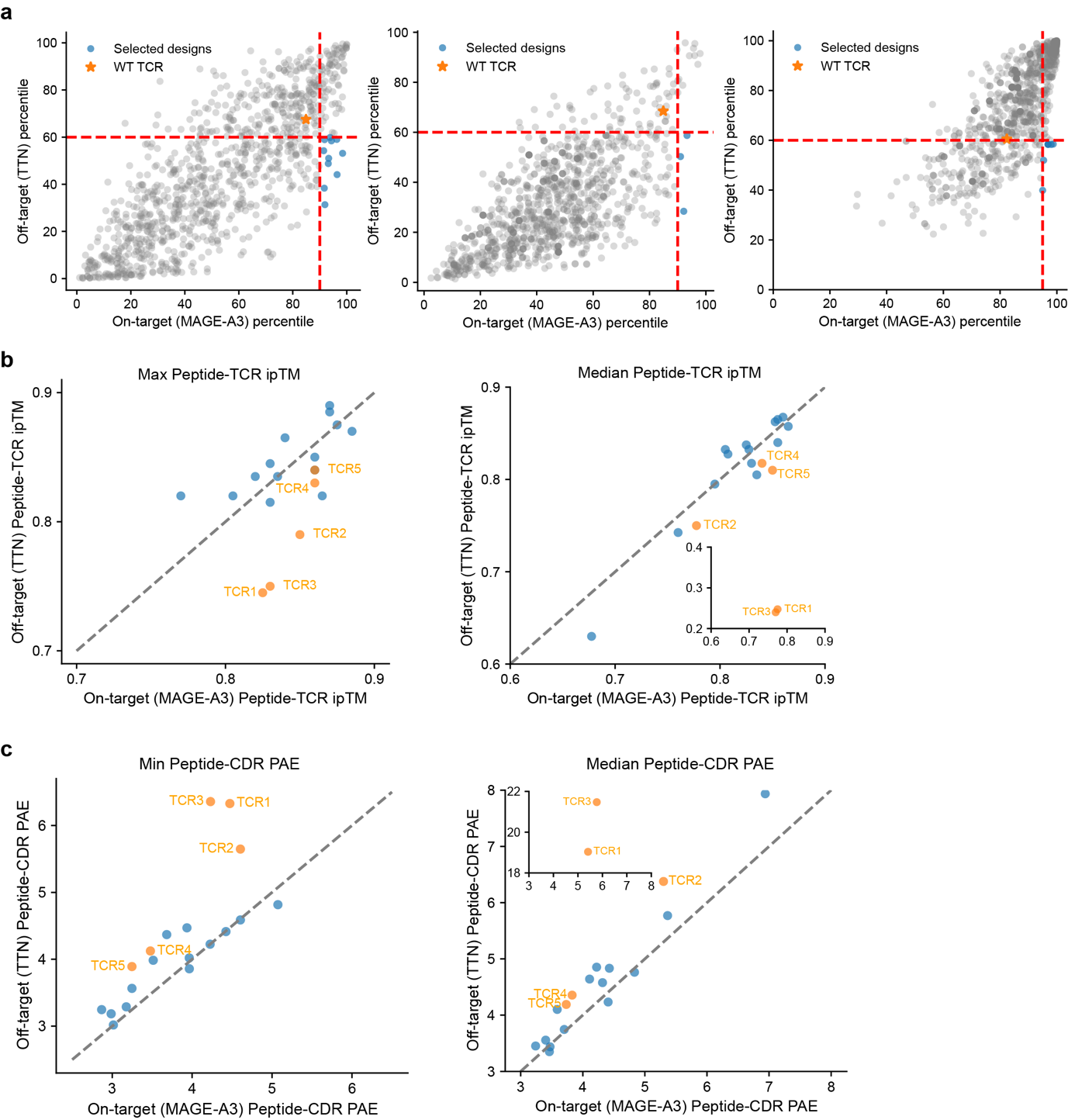

**Supplementary Fig. 10. Filtering of MAGE-A3 antigen-specific TCR candidates using binding and structure prediction.** **a,** The distributions of on-target (MAGE-A3) and off-target (TTN) binding rank percentiles generated by (left) TCRDiff, (middle) TCRDiff+, and TCRDiff (beta-only). Dash lines, on-target/off-target percentile thresholds. **b,** Predicted on-target (MAGE-A3) and off-target (TTN) peptide-TCR ipTM of the candidate TCR designs. **c,** Predicted on-target (MAGE-A3) and off-target (TTN) peptide-CDR PAE of the candidate TCR designs.

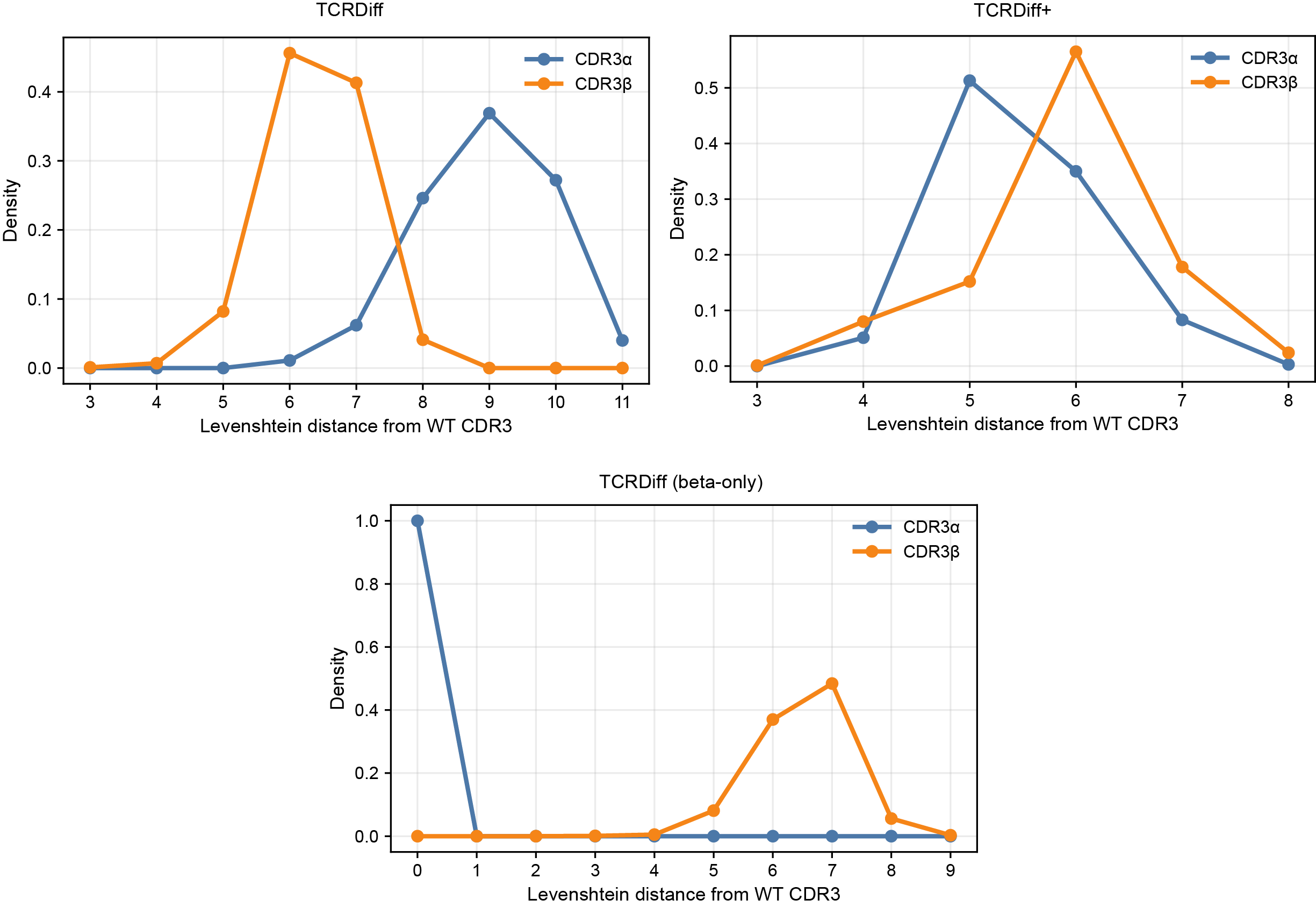

**Supplementary Fig. 11. Distance distribution of the generated CDR3αβ sequence from the wild-type MAG-IC3 TCR.** Density plots of the Levenshtein distances between generated CDR3αβ sequences and wild-type CDR3. The generated sequences were derived from (top left) TCRDiff, (top right) TCRDiff+, and (bottom) TCRDiff (beta-only).

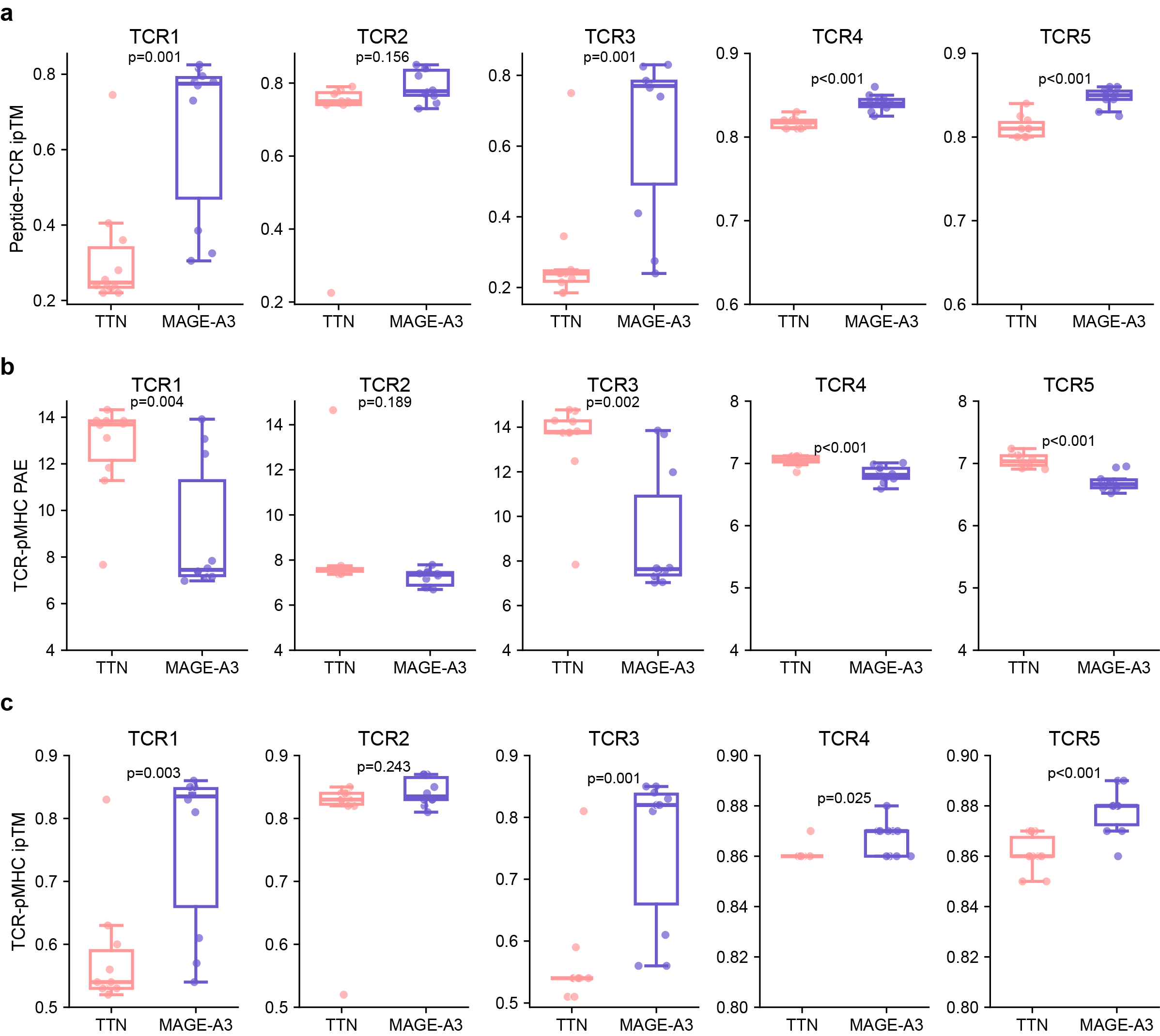

**Supplementary Fig. 12. Confidence metrics of the predicted TCR-pMHC complex structures.** Additional structural confidence metrics of predicted TCR-pMHC complexes for five candidate TCR designs under ten random seeds. **a,** peptide-TCR ipTM; **b,** mean TCR-pMHC PAE; **c,** TCR-pMHC ipTM. P-values were determined by Student’s t-test. Box center line, median; box limits, upper and lower quartiles; whiskers, 1.5 × interquartile range; points, data points.
